# Response of the organellar and nuclear (post)transcriptomes of Arabidopsis to drought stress

**DOI:** 10.1101/2022.08.09.503311

**Authors:** Duorong Xu, Qian Tang, Dario Leister, Tatjana Kleine

## Abstract

Plants have evolved sophisticated mechanisms to cope with drought, which involve massive changes in nuclear gene expression. However, little is known about the roles of post-transcriptional processing of nuclear or organellar transcripts and how meaningful these changes are. To address these issues, we used long non-coding RNA-sequencing to monitor (post)transcriptional changes during different times of drought exposure in Arabidopsis Col-0 and a mutant (*protein phosphatase 7-like*, *pp7l*), from which we demonstrated that it can survive long periods of drought stress. The changes detected in the *pp7l* mutant were marginal, while in the wild type chloroplast transcript levels were globally reduced, editing efficiency dropped, but splicing was not affected. Mitochondrial transcripts were slightly elevated, while editing and splicing were unchanged. Also, transcriptional activation of transposable elements played only a minor role. Conversely, alternative splicing (AS) affected nearly 2,000 genes (11% of expressed nuclear genes). Of these, 25% underwent isoform switching, and 15% were regulated solely at the level of AS, representing transcripts that would have gone unnoticed in a microarray-based approach. Our data show that AS enhances proteome diversity to counteract drought stress and represent a valuable resource that will facilitate the development of new strategies to improve plant performance under drought. Moreover, altering the relative contributions of spliced isoforms might enhance drought resistance. For instance, our data imply that accumulation of a non-functional *FLM* (*FLOWERING LOCUS M*) isoform – and not the ratio of functional isoforms as suggested for temperature responses - accounts for the early-flowering phenotype under drought conditions.

## INTRODUCTION

Land plants must cope with all sorts of environmental conditions, since, as sessile organisms, they cannot evade them. Owing to climate change, the frequency and amplitude of extreme conditions are increasing, and this seriously threatens crop yields worldwide (Zhao et al., 2017). Drought is the most important abiotic stress (Singh et al., 2018), and responses to drought are influenced by developmental stage, plant species and degree of stress (Hu and Xiong, 2014). The primary strain during drought stress is dehydration – loss of water from the cell – which triggers osmotic and hormone-related signals (Blum, 2016), in particular the phytohormone abscisic acid (ABA) (Zhu, 2002). Terminology and approaches to drought research vary widely and the issues are often oversimplified (Lawlor, 2013; Blum and Tuberosa, 2018; Kooyers, 2019). Taking our cue from the work of Lawlor (2013) and Blum (2016) and the TNAU Agritech Portal (https://agritech.tnau.ac.in/agriculture/agri_drought_tolerent_mechanism.html), we define drought resistance in terms of the following features: (i) drought survival, ii) drought escape, iii) drought avoidance, and iv) drought tolerance. Drought survival refers to the fact that, under drought conditions, cells, tissues, and organs are able to maintain key cellular functions and can recover, with minimal damage, upon relief of drought stress (Lawlor, 2013). Dehydration avoidance is a response to moderate, temporary drought stress, which involves deposition of cuticular waxes and growth retardation, in addition to the reduction of transpiration via ABA-mediated closure of stomata (Gupta et al., 2020). In the drought-escape strategy, which is employed by annual plants like *Arabidopsis thaliana* (Arabidopsis), flowering is accelerated before drought can compromise survival of the plant (Kenney et al., 2014). In drought tolerance, the plant is able to maintain its functions under dehydration – for example, by producing larger amounts of sugars, osmoprotectants, antioxidants, and scavengers of reactive oxygen species (Hu and Xiong, 2014).

Responses to drought and other stresses are accompanied by large-scale changes in gene expression that facilitate the production of compounds needed to counteract the imposed strain (Bashir et al., 2019). Thus, transcriptome-based studies have identified acetate as a compound that helps plants to survive drought stress (Kim et al., 2017), and shown that the transcription factor AGAMOUS-LIKE 22 (also known as SHORT VEGETATIVE PHASE, SVP), which participates in primary metabolism and developmental processes, is activated in drought-stressed Arabidopsis plants (Bechtold et al., 2016).

Changes in nuclear gene expression in response to environmental perturbations, including drought stress, are fine-tuned by retrograde signals from the chloroplast (Estavillo et al., 2011; Kleine and Leister, 2016), and chloroplasts act both as a target and as a sensor of environmental changes (Kleine et al., 2021). In a forward genetic screen two chloroplast proteins involved in drought resistance were identified (Hong et al., 2020). Furthermore, the first steps of ABA biosynthesis take place in plastids (Ali et al., 2020), which underlines the importance of the chloroplast in the response to drought stress. Although chloroplasts and mitochondria each harbor only around 100 genes (Kleine et al., 2009), organellar gene expression is complex, and transcripts can undergo post-transcriptional modification events such as splicing and editing (Germain et al., 2013). Therefore, the abundance and functionality of a number of organellar proteins depends not only on the levels of their transcripts, but also on their modification status. The same holds true for nucleus-encoded transcripts: Here, alternative splicing (AS) provides for the synthesis of different transcript isoforms from the same gene, thereby increasing proteome diversity (Laloum et al., 2018; Bashir et al., 2019). It has become clear that AS is of central importance for abiotic stress tolerance in plants (Laloum et al., 2018), and AS itself is dynamically regulated under cold stress (Calixto et al., 2018). Concerning organellar transcriptome changes, it has been demonstrated that heat stress over periods of several hours increases the abundance of chloroplast transcripts, and induces a global reduction in splicing and editing efficiency (Castandet et al., 2016), but to our knowledge, the behavior of the mitochondrial (post)transcriptomic landscape has not been investigated so far.

However, microarray analysis has been the major source of information on transcriptome changes in response to drought (Bechtold et al., 2016; Kim et al., 2017; Bashir et al., 2019), while transcriptome-wide AS analysis in response to cold has been addressed with the aid of mRNA sequencing (mRNA-Seq) (Calixto et al., 2018). Both methods make use of samples enriched for polyadenylated transcripts, and are designed specifically to trace the accumulation of nuclear transcripts. Functional organellar transcripts are not polyadenylated; indeed, unlike nuclear transcripts, they become unstable upon addition of a poly(A) tail (Rorbach et al., 2014).

Therefore, we set out to extend the investigation of changes in nuclear gene expression under drought stress by undertaking an overarching characterization of the post(transcriptome), including alternative splicing of nuclear transcripts, accumulation of organellar (chloroplast and mitochondrion) transcripts, editing and splicing of organellar transcripts, and the transcriptional activation of transposable elements. To this end, long non-coding (lnc) RNA-Seq was applied, because its workflow involves library preparation after depletion of ribosomal RNA instead of enrichment for mRNAs as in mRNA-Seq. For reproducibility, an already published drought kinetics set-up (Kim et al., 2017) was applied. Moreover, we compared the transcriptional behavior of the wild type (Col-0) to that of the *pp7l* mutant, which is devoid of PP7L, a member of a small subfamily of phosphoprotein phosphatases (Xu et al., 2019). We chose this mutant because, as we show here, it can survive very long periods of drought stress (in contrast to Col- 0). Moreover, based on our platform, we describe changes in the various organellar and nuclear (post)transcriptomes under drought and complement this with an analysis of transcript isoform switches. In particular, we suggest an FLM (FLOWERING LOCUS M)-dependent early flowering mechanism under drought, which is characterized by massive production of non-functional *FLM* isoforms.

## RESULTS

### The *pp7l* mutant can survive long periods of drought stress – and will be used together with Col-0 in RNA-Seq experiments

To investigate the impact of drought stress on the (post)transcriptome, we wanted to include for comparison a mutant that can cope better than wild type with drought stress. In *Arabidopsis thaliana*, loss of PP7L, a member of a small subfamily of phosphoprotein phosphatases (PPPs), results in pleiotropic phenotypes including decreased resistance to salt, high-light and temperature stress (Xu et al., 2019; Xu et al., 2020). During further phenotypical characterization, we noticed that *pp7l* mutants also displayed better tolerance of drought stress after water was withheld for 15 days from 3-week-old Col-0 and *pp7l* mutant plants. Interestingly, upon re-watering of these plants for three days, 60% of them survived, while all Col-0 plants died (Supplemental Figure S1). However, *pp7l* is a dwarf mutant (Xu et al., 2019). Thus, it is possible that *pp7l* and other dwarf mutants are more resistant to drought conditions because their roots extract water from the soil at a lower rate. To investigate this in more depth, we performed a second drought experiment in which we exposed Col-0 and *pp7l* to the same drought conditions by controlled watering through adjusting pot weights (Figure 1A). Remarkably, Col-0 plants died after 46 days of drought treatment while the *pp7l* mutant still displayed photosynthetic activity as shown by the measurement of maximum quantum yield of photosystem (PS) II (Fv/Fm; Figure 1A). In another experimental set-up, we included the dwarf mutants *cop1* (Moazzam-Jazi et al., 2018) and *sal1* (Wilson et al., 2009) which have been described before as drought resistant as controls. Because COP1 likely acts downstream of the cryptochrome (CRY) pathway as a repressor of stomatal opening, and non-dwarf *cry1 cry2* mutants are also drought tolerant (Mao et al., 2005), this double mutant was also included in our study. Water was withheld from 3-week-old plants and pictures were taken after 16 and 24 days. All mutants appeared to be drought tolerant after 16 days. They accumulated less anthocyanin and leaves looked healthier than those of the WT (Figure 1B). However, the remarkable ability of the *pp7l* mutant to cope with drought stress was underpinned after 24 days. After this period, *pp7l* showed no signs of wilting, while *sal1* had begun to wilt, and *cop1- 4* and *cry1 cry2* were clearly wilting.

**Figure 1.**
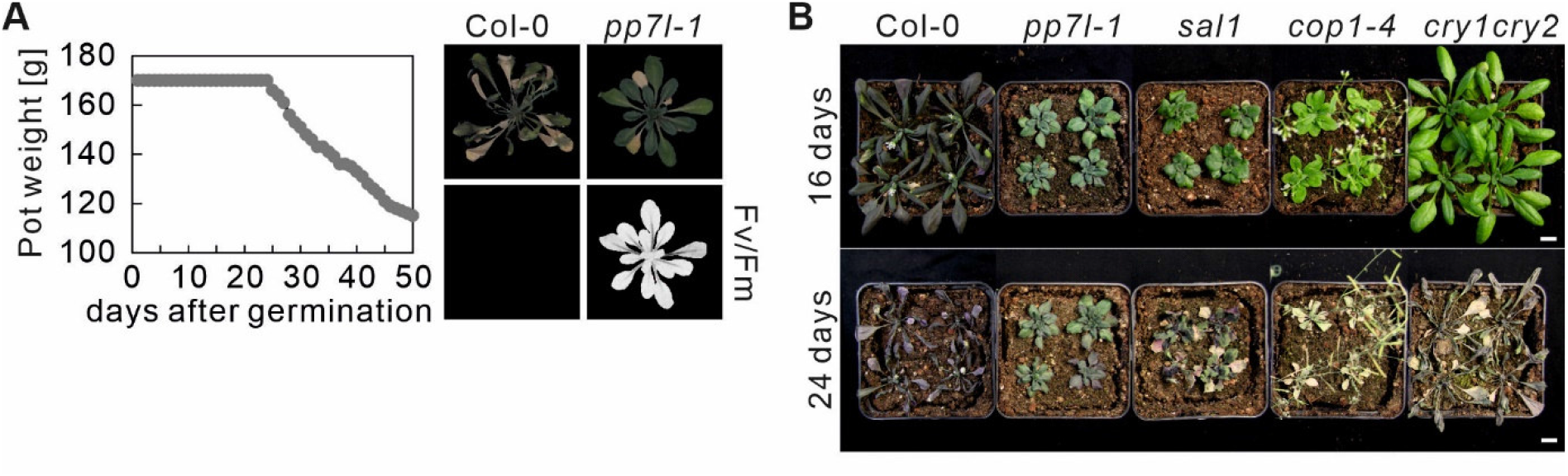
Soil-grown *pp7l* plants are more drought tolerant than the wild type. **A,** The water status was controlled by watering pots to the same weight. After 24 days pot weight was gradually reduced. 46-day-old Col-0 plants died while *pp7l* plants were still viable as indicated by detectable photosynthetic activity (maximum quantum yield of photosystem II, Fv/Fm). **B,** Col-0 and the different genotypes were grown for 3 weeks under control conditions, and then water was withheld for 16 and 24 days. Bar = 1 cm.

In conclusion, the hypothesis of differential water usage by the dwarf mutants cannot be dismissed, however we can classify *pp7l* as a bona fide survivor of long drought periods.

### Drought-related changes in gene expression are robust across different laboratories

Our goal was to investigate in addition to differential gene expression, the behavior of post-transcriptional processing of nuclear or organellar transcripts under drought. To this end, long non-coding RNA sequencing (lncRNA-Seq) was performed with RNA isolated from 3-week-old Col-0, but also *pp7l* plants (prompted by the long survival under drought stress) grown under optimal conditions (time-point 0 days; 0d_DS), and after withholding of water for 6, 9, 12 and 15 days (6d_DS, 9d_DS, 12d_DS and 15d_DS, respectively) as described previously (Kim et al., 2017). Here it is of note, that we deliberately chose the “water-withheld-setup”, instead of a strictly controlled one because of reasons stated in the Discussion section. As a second note, Kim et al. (2017) used microarrays, which are not suitable for our purpose – monitoring of organellar transcript accumulation and editing events, and the investigation of splicing of transcripts produced in organelles as well as the nucleus. We repeated our experiment separately with different batches of plants, yielding three biological repeats per time point, except for Col-0 6d_DS (two replicates). The sequencing depth was approximately 25 million 150-bp paired-end reads for each of the samples (Supplemental Table S1).

Investigation of whole-genome wide gene expression changes showed that 16,850 genes were considered as expressed after removal of weakly expressed ones, and most of them (14,173) were differentially expressed in response to drought when compared with the starting condition or between genotypes (Figure 2A, Supplemental Table S2). These findings show that drought stress has a substantial impact on the transcriptome. Here, a gene was considered to be differentially expressed (DEG) if it showed an absolute log_2_ fold change ≥1 (≥ 2-fold linear change) in expression in at least one contrast group (adjusted *P* < 0.01). In Col-0, 49.2% were down- and 50.8% upregulated, respectively, while in *pp7l* slightly more genes were down- (54.4%) than upregulated, but the differences in gene expression increased only moderately (Figure 2A). This is also reflected in a principal-component analysis (PCA) of gene expression data from across the 29 samples, which showed that drought (64% of variance) was the major driver of gene expression in Col-0, but not in *pp7l* (Figure 2B).

**Figure 2.**
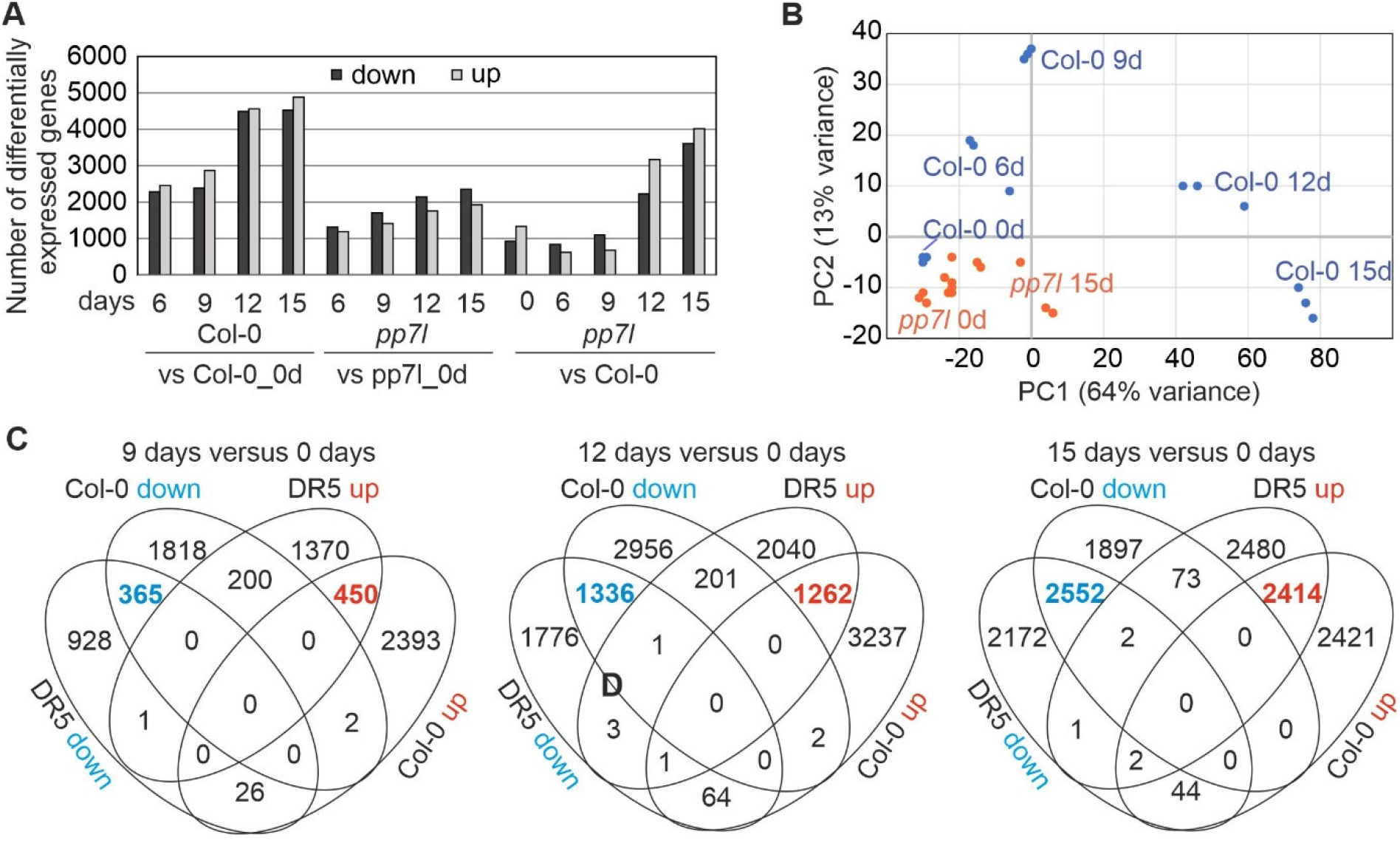
Drought-related changes in gene expression are robust across different laboratories. **A**, Numbers of differentially expressed genes (absolute log_2_ fold change ≥1; in at least one contrast group with an adjusted *P* < 0.01) in Col-0 and *pp7l* plants subjected to drought exposure compared to the respective control (0 days), and comparisons of *pp7l* to Col-0 at the indicated time points. **B,** Principal Component Analysis (PCA) plot visualizing variation between genotypes and time-points based on RNA-seq data. **C,** Comparison of transcript changes evoked by drought stress in Col-0 (this study) or in DR5 (Kim et al. 2017). Venn diagrams illustrate the total numbers of differentially expressed genes shared between or specific for the different treatments.

Our re-analysis of data published by Kim et al. (2017), and comparison with the transcriptome data generated in the present study, showed that drought-related changes in gene expression are robust across different laboratories, wild-types (Col-0 and DR5, a Col-0 line containing a *DR5*::*GUS* construct), transcriptome platforms, and sampling methods (whole plants in our set-up vs. aerial parts in the work of Kim et al.) after prolonged exposure to drought stress (Figure 2C). Accordingly, we provide lists of robust drought DEGs in Supplemental Table S3. Some of these were exceedingly differentially expressed. For example, levels of *ALDEHYDE DEHYDROGENASE 7B4* (*ALDH7B4*) mRNA were 4-fold higher after 9 days of drought treatment, but had risen by more than 3000-fold after 12 and 15 days.

### Gene ontology analysis

The kinetic data were further explored by sorting the DEGs into nine different clusters, across genotypes and duration of drought stress. Cluster 1 contained the genes that were strongly downregulated in Col-0 at 12d_DS and 15d_DS, but only moderately repressed in *pp7l*. Gene Ontology (GO) analysis of this cluster showed enrichments of genes encoding proteins related to the response to low light, protein folding, chlorophyll biosynthesis, cp translation and photosynthesis-related processes in the biological process (BP) category (Figure 3A and B, Supplemental Table S4). This is in line with the phenotypes and Fv/Fm values of Col-0 and *pp7l* under drought stress (see Figure 1) and underlines the greater susceptibility of Col-0 to drought stress. Moreover, the GO analysis revealed that, in addition to cp-encoded genes (see Figure 6), nucleus-encoded chloroplast proteins are also regulated at the transcript level. In the cellular component (CC) category, transcripts coding for components of the PSI, PSII and the cp Ndh complexes, as well as cp and cytosolic ribosomal subunits, were enriched. The similarities in the behavior of transcripts for both cp and cytosolic ribosomal subunits reflect the importance of coordination between protein synthesis in chloroplasts and the cytosol (Wang et al., 2018) during drought stress. The majority of mt transcripts in Col-0 increased or barely changed under drought stress (see Supplemental Figure S6). Accordingly, genes encoding mt proteins were found in cluster 6, which encompasses genes whose transcript levels especially increased in Col-0 at 9d_DS and 12d_DS (Figure 3), in particular those encoding proteins of the substrate-carrier family, succinate-dehydrogenase complex II and enzymes of the tricarboxylic acid cycle (TCA). Indeed, TCA cycle metabolites are known to increase after water shortage (Pires et al., 2016). Cluster 6 also includes the categories “peroxisome”, “response to water” and “response to osmotic stress”. The latter two categories both point to a disturbance in water balance, and genes encoding proteins involved in responses to hypoxia, abscisic acid and salt stress are found in cluster 5, which encompasses genes that behave like those of cluster 6. Moreover, genes assigned to the GO categories “cellular water homeostasis”, together with “cell wall thickening” and “fatty acid biosynthesis” in cluster 2 are specifically induced in Col-0 after 6 or 9 days of drought stress.

**Figure 3.**
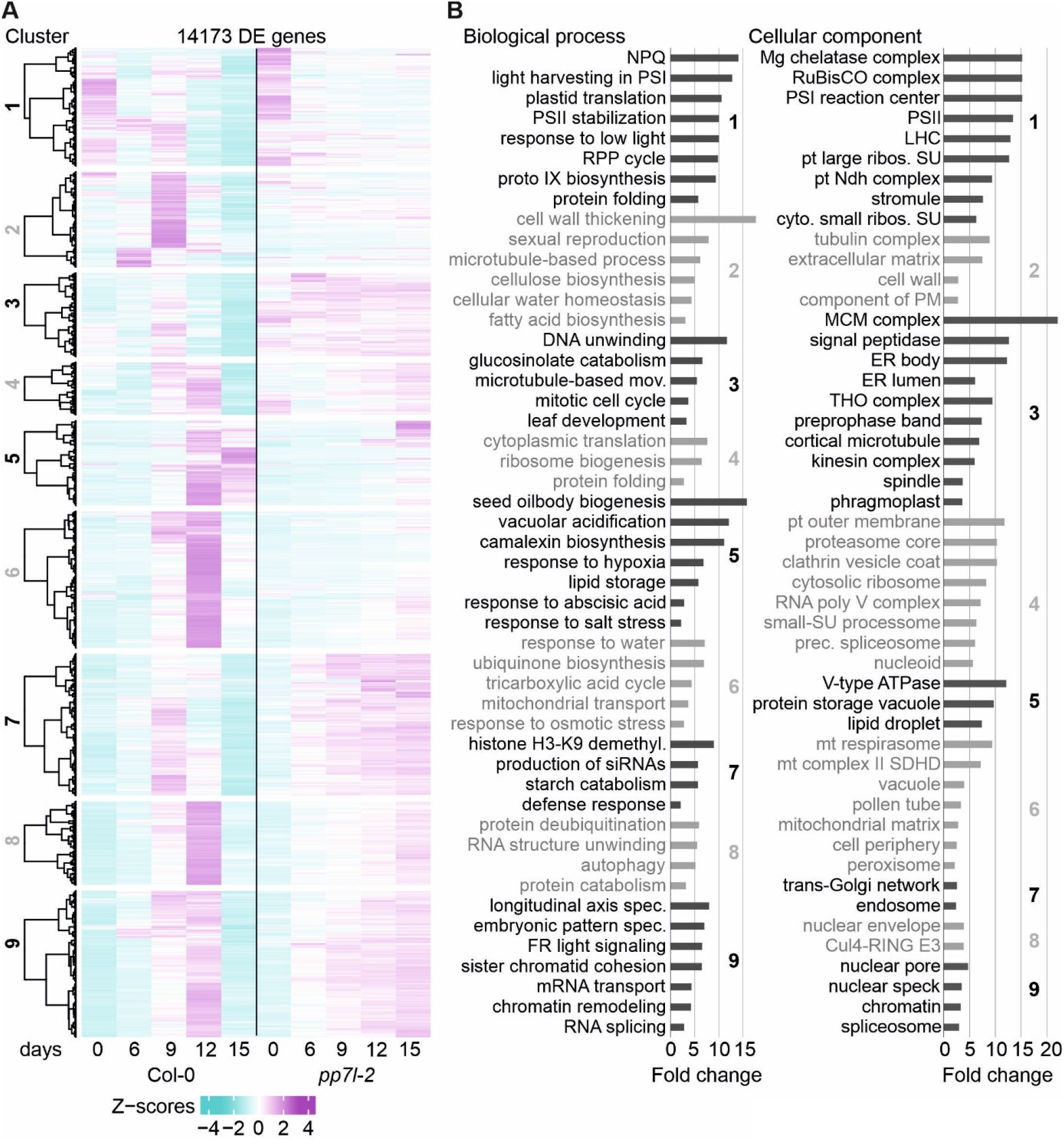
Overview and gene ontology analysis of gene expression changes under drought. **A,** Heatmap of differentially expressed (DE) genes under drought stress compared to the control time point (0 days). Hierarchical clustering was used to partition the DE genes into nine clusters with the Euclidean distance and ward.D clustering algorithm. **B,** Graphs illustrating non- redundant Gene Ontology (GO) term enrichment for the biological process and cellular component categories according to DAVID (Huang da et al., 2009) and REVIGO (Supek et al., 2011). GO terms with a >2-fold change and a Benjamini-corrected *P*-value of <0.05 are shown. Cul4-RING E3, Cul4-RING E3 ubiquitin ligase; LHC, light-harvesting complex; mov., movement; NPQ, nonphotochemical quenching; prec., precatalytic; protein catabolism, ubiquitin-dependent protein catabolism; RPP, reductive pentose-phosphate; spec., specification; PS, photosystem; pt, plastid; ribos., ribosomal; SU, subunit.

In summary, drought stress induces massive transcriptional reprogramming, and the polarities of changes in gene expression (up or down) are compatible with responses to drought at the phenotypical and metabolic levels. In addition, coordination of the organellar and nuclear genomes at the transcript level is apparent.

### Transcription of transposable elements is not released under drought stress

Cluster 7, which is characterized by the transient up-regulation of specific mRNAs in Col-0 at 9d_DS and is constitutively up-regulated in *pp7l* at all stages of drought stress, attracted our attention. Besides “starch catabolism”, “histone H3-K9 demethylation” and “production of siRNAs”, it contains the category “defense response”. This last group mainly comprises disease resistance proteins of various classes, along with the transcripts encoding the photoreceptors cryptochrome 1 (cry1) and cry2, which have previously been implicated in the drought-resistance response (see the control in Fig. 1, and Mao et al., 2005). The category “histone H3- K9 demethylation” is represented in cluster 7 by several jumonji (jmjC)-domain-containing proteins, including JMJ27, which positively regulates drought-stress responses by mediating histone H3K9 demethylation (Wang et al., 2021), confirming the ability of our approach to identify new players in drought stress.

This finding prompted us to explore the putative involvement of the RNA-directed DNA methylation (RdDM) pathway in the response to drought treatment, we focused on the genes listed in “production of siRNAs” category. These genes code for NRPD1A and NRPD2A (subunits of the nuclear RNA polymerase IV), the RNA-dependent RNA polymerases RDR2 and RDR6, DICER-LIKE 3 (DCL3), ARGONAUTE 4 (AGO4), and DEFECTIVE IN RNA-DIRECTED DNA METHYLATION 1 (DRD1). With the exception of RDR6, these proteins are all involved in the RdDM pathway (Wendte and Pikaard, 2017) in which 23- and 24- nucleotide (nt) small interfering RNAs (siRNAs) (Singh et al., 2019) are produced and loaded mainly onto AGO4 and AGO6. Ultimately, an AGO-siRNA-Pol V complex recruits chromatin-modifying enzymes that catalyze *de-novo* DNA methylation to silence invasive nucleic acids of viral, transposon or transgenic origin, as well as endogenous genes (Wendte and Pikaard, 2017; Zhang et al., 2018). Moreover, RdDM participates in responses to abiotic stress (Liu and He, 2020), and loss of RDR2, DCL3 or AGO4 decreases thermotolerance (Popova et al., 2013). Therefore, we investigated the expression of transposable elements in response to drought stress. At first sight, it appeared that nearly 300 TEs were indeed transcriptionally activated specifically in Col-0 after prolonged drought exposure (Figure 4A, Supplemental Table S5). However, closer inspection of the read distribution across the TEs, and overlaps between the corresponding TE genome intervals with the genome positions of protein-coding genes, showed that only a minor fraction could be attributed to bona fide TEs (Figure 4B; Supplemental Figure S2). Other instances of up-regulation were detected in the intronic regions of TEs or coincided with up-regulated transcripts that overlapped with TEs (Supplemental Figure S2).

**Figure 4.**
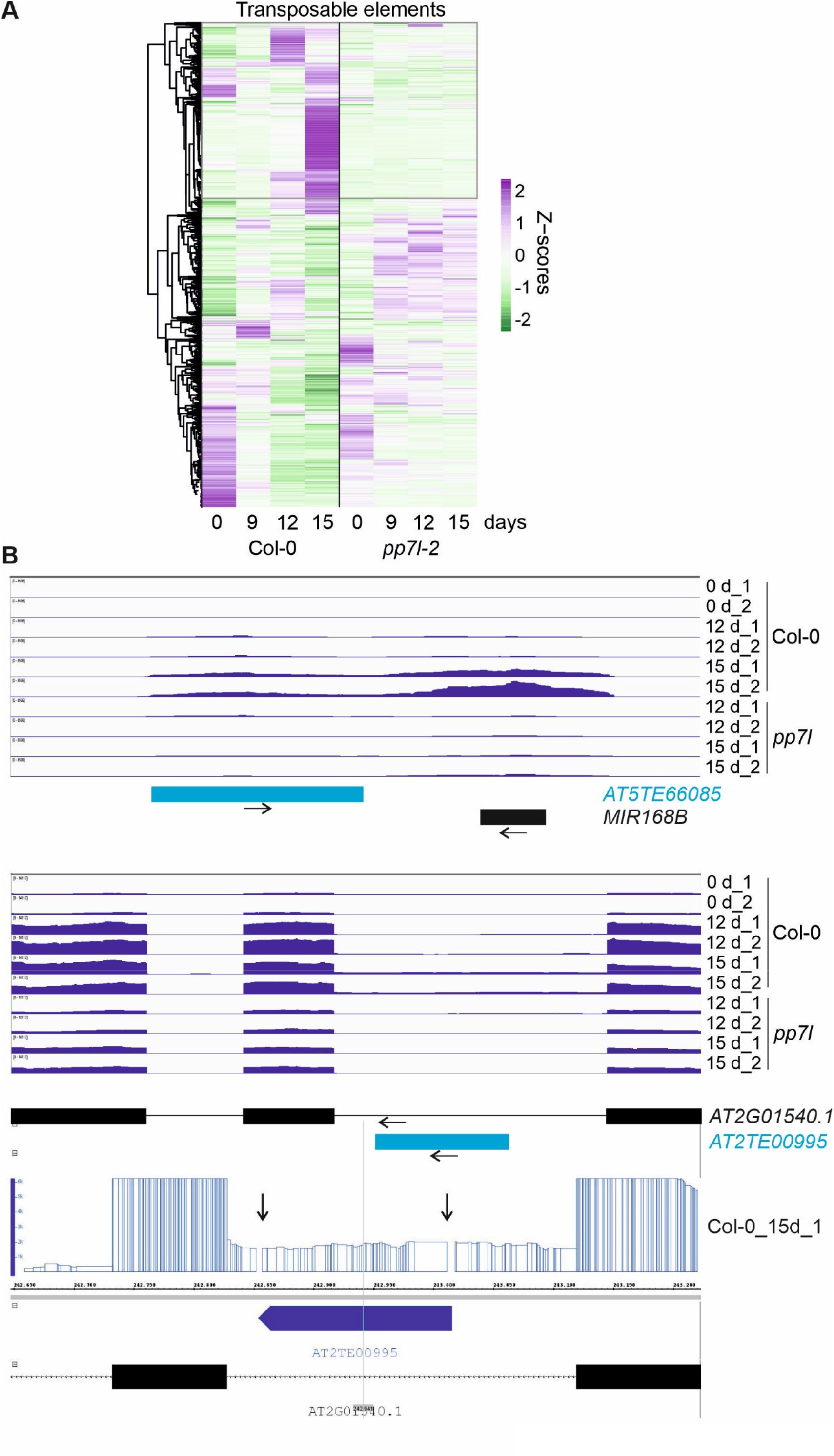
Transcriptional activity of transposable elements (TEs) under drought stress. **A,** Heatmap illustrating accumulation of TE transcripts (displayed as Z-scores) under control conditions (0 days) and after 9, 12 and 15 days of drought stress. Low to high expression is represented by green to purple transition. Note that Z-scores are calculated for each individual TE over the time-course. **B,** Pattern of transcript accumulation covering regions of TEs. The normalized read depths of transcripts detected in Col-0 and *pp7l* plants under control and prolonged drought conditions were visualized with the Integrative Genomics Viewer (IGV). Transposable elements (blue boxes), exons (black boxes), introns (black lines) and the 5′- and 3′ UTRs (gray boxes) are shown.

Hence, it can be concluded that transcription of transposable elements is not widely activated under drought stress.

### Drought stress has a negative impact on the chloroplast (post)transcriptome

Heat stress for periods of several hours induces a global reduction in splicing and editing efficiency in chloroplasts (cp), and an overall increase in the abundance of cp transcripts, while short-term drought stress for 3 and 12 hours results in only minor changes in cp transcripts (Castandet et al., 2016).

To investigate cp transcripts under drought stress, reads were mapped and processed with the help of ChloroSeq (Castandet et al., 2016). Overall levels of noncoding cp RNAs are higher after 12 h of heat treatment (Castandet et al., 2016), but inspection of our bamcoverage files, which include reads across all nucleotide (nt) positions of the cp genome, showed that this does not hold for exposure to drought (Supplemental Figure S3). In fact, cp coverage already indicated lower overall cp transcript accumulation in Col-0 relative to *pp7l* after prolonged drought stress (Supplemental Figure S3). To investigate this for individual transcripts, heatmaps of z-means (Figure 5A) and fold changes of cp transcripts (Supplemental Table S6A) were generated, which revealed substantial reductions in levels of most transcripts under prolonged drought conditions in Col-0, while the general trend in the *pp7l* mutant pointed in the opposite direction (Figure 5A, Supplemental Table S6A). Notably, in 4-day-old *pp7l* seedlings grown under well-watered conditions, cp transcripts also accumulated to higher levels than those observed in wild-type controls (Xu et al., 2019). Here, it should be noted that analysis of tRNAs was excluded (also in the following editing and splice-site analyses) because their small size and many modifications make them difficult to amplify with either the lncRNA-Seq method or conventional mRNA-Seq protocols. Of the 69 protein-coding cp genes that were above the detection threshold, 50, 63 and 64 were at least 2-fold reduced (compared to the 0d_DS sample) in Col-0 plants after 9, 12 and 15 days of drought, respectively. The genes most affected (those whose transcripts were reduced to 3% or less of their initial levels) code for the D3, F and K subunits of the NAD(P)H dehydrogenase complex and a protein of the large ribosomal subunit (rpl23). In *pp7l* these transcripts were still present at 30-60% of their initial levels after 15d_DS. The relatively constant – and transiently even higher – expression of genes encoding the PSII reaction-center proteins PsbA, PsbC and PsbD in *pp7l* plants throughout drought stress is particularly noteworthy (Figure 5A, Supplemental Table S6A).

**Figure 5.**
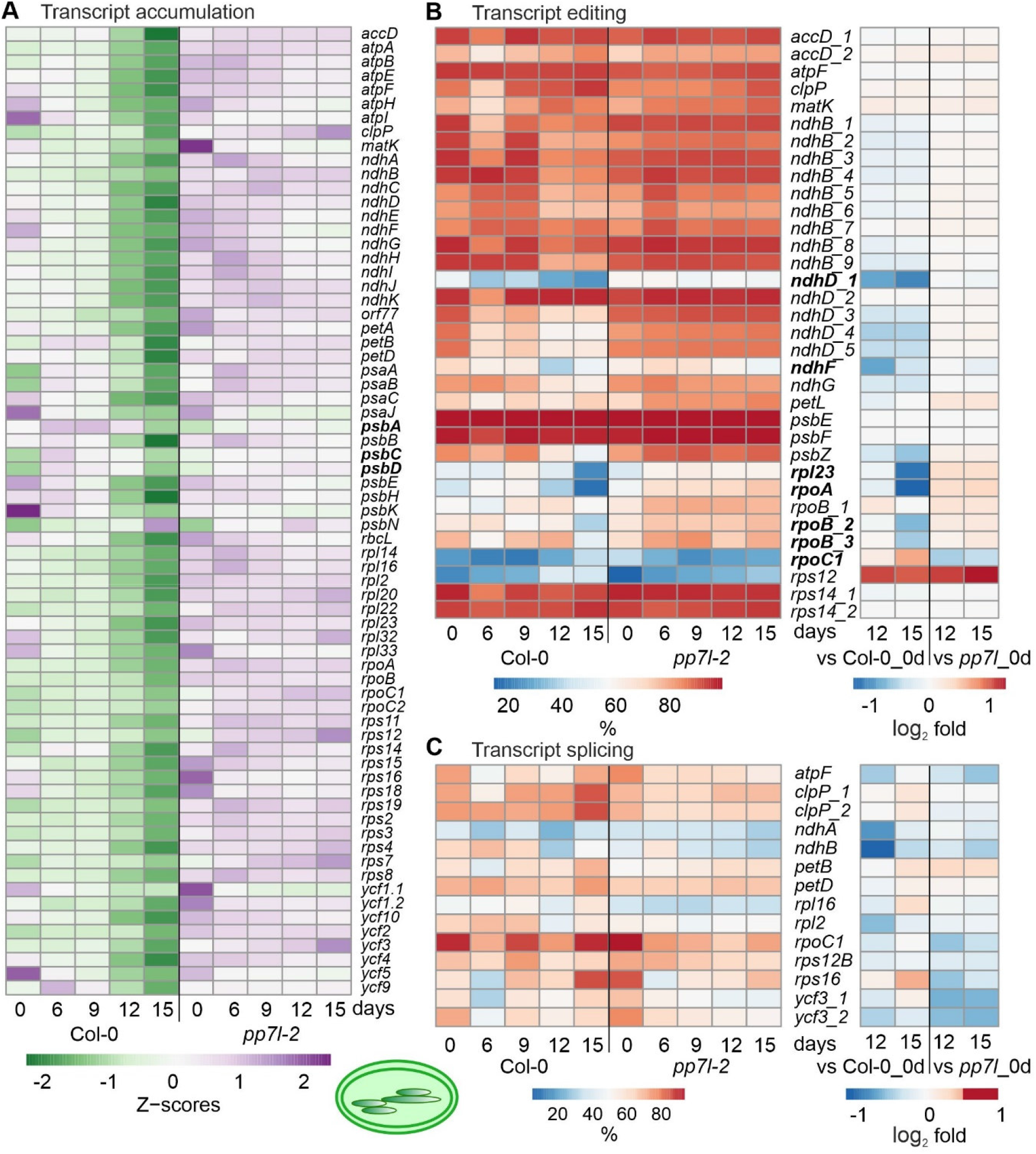
Impact of drought stress on accumulation, editing and splicing of chloroplast transcripts. Water was withheld for 15 days from 3-week-old Col-0 and *pp7l* plants grown under standard conditions (0 days), and lncRNA-Seq was performed as described in Materials and Methods on RNA extracted from plants harvested at the indicated time-points. **A,** Heatmap illustrating chloroplast transcript accumulation (Z-scores) during the drought time-course. Low to high expression is represented by the green to purple transition. Note that Z-scores are calculated for each individual transcript over the time course. **B, C,** Percentages of editing **(B)** and splicing (C) events during the time-course (left) and log_2_ fold changes after 12 and 15 days of drought stress compared to the control time point (right).

To examine post-transcriptional changes in cp transcripts, editing and splicing efficiencies were calculated. Interestingly, unlike transcript accumulation, editing was not generally reduced in Col-0 under drought conditions, but was observed especially for *ndhD*, *ndhF*, *rpl23* and the *rpoA* and *rpoB* genes encoding α and β subunits of plastid-encoded RNA polymerase (Figure 5B). Moreover, editing of *rpoC1* was slightly enhanced after 15d in Col-0, but reduced in *pp7l*. Editing of *rps12* was enhanced in both Col-0 and *pp7l*, which suggests that editing of this transcript is not a drought-specific effect. After 15 days of drought stress, splicing efficiency was not notably diminished, and in some cases (five out of 14 events) was slightly enhanced (Figure 5C).

To summarize, while overall cp mRNA levels were broadly reduced in the WT, impairment of editing events was more marked, and splicing was not compromised at all. Moreover, the (post)transcriptomic landscape of *pp7l* under drought was only slightly altered.

### Mitochondrial transcript accumulation, editing and splicing under drought

Mitochondrial (mt) transcripts were also investigated, and inspection of bamcoverage files of the mt genome suggested even higher mt transcript accumulation under drought stress for Col- 0 (Supplemental Figure S4), in contrast to the fall in levels of cp transcripts (see Supplemental Figure S3). This was confirmed by producing heatmaps of z-means of mt transcripts and their fold changes (Supplemental Figure S5A, Supplemental Table S6B). Thus, levels of the majority of mt mRNAs in Col-0 increased or were only slightly changed during drought stress, while the *pp7l* mutant was again significantly less affected (Supplemental Figure S5A, Supplemental Table S6B). Of the 49 protein-coding mt genes that were above the detection threshold, 11, 11 and 26 transcripts were at least 2-fold elevated in Col-0 after 9, 12 and 15 days of drought, respectively, compared to 0d_DS. Only a few mRNAs were down-regulated during drought stress, of which the two most reduced transcripts *ATP synthase subunit 1* (*atp1*; 0.3%) and *rpsL2* (17.8%) were detected at 15d_DS. Interestingly, transcripts coding for mt NADH dehydrogenase subunits increased during prolonged drought stress – in stark contrast to their counterparts in the cp sister complex. In *pp7l*, mt *nad* transcripts were only moderately increased, if at all.

With regard to post-transcriptional changes of mt transcripts in Col-0, transient down- regulation of splicing efficiency was observed for *cytochrome oxidase 2* (*cox2*), *nad1* and *nad7- 2*, and splicing capacity was slightly enhanced after 15 days (Supplemental Figure S5B). Editing capacity under drought was not changed relative to 0d_DS, except in the case of *ccb206* (encoding the cytochrome *c* biogenesis protein 206), which was reduced in Col-0 at 12d_DS (Supplemental Figure 6C). But this change was also seen in *pp7l*, therefore, this might presumably not be a drought specific effect.

Overall therefore, mt (post)transcription was not markedly impaired in either Col-0 or *pp7l* and drought-induced changes tended to show the opposite trend to that seen in the cp transcripts. However, the drastic reduction of *atp1* transcript levels in Col-0 plants after 15 days of drought stress is particularly striking.

### Splicing of nuclear transcripts under drought stress

Coverage plots at the loci *AT2TE82150* and *AT1TE17050* (see Supplemental Figure S2) suggested differential isoform usage of *AT2G43780* and intron retention in *AT1G15340*, respectively, after drought stress. Indeed, DAVID detected an enrichment of the GOs “RNA splicing/spliceosome” and “the pre-catalytic spliceosome” in clusters 9 and 4, respectively, which host genes whose transcripts were transiently up-regulated in Col-0, but remained high in *pp7l* (see Figure 3). To systematically investigate splicing behavior under drought stress, genome-wide differential alternative splicing (DAS) and differential transcript usage (DTU) during drought stress was examined with the help of the high-quality AtRTD2_QUALI transcriptome and 3D RNA-Seq (Zhang et al., 2017; Guo et al., 2021). By this means, in addition to the expression at the gene level (i.e., the sum of all transcript abundances of a given gene), the individual transcript isoform levels could be determined. In all, 1,988 DAS genes were identified, of which 1,680 were also DE genes (regulated by both transcription and AS), while 308 genes were regulated by AS only in at least one contrast group (Figure 6A, Supplemental Table S7). Among the latter are LESION SIMULATING DISEASE 1 (LSD1), which monitors a superoxide-dependent signal and negatively regulates plant cell death (Bernacki et al., 2021), as well as playing an important role in survival of drought stress (Szechynska-Hebda et al., 2016), plastidic acetyl-CoA synthetase (ACS), REGULATORY PARTICLE TRIPLE-A ATPASE 6A (RPT6A) (found by Co-IP) which is a component of the 26S proteasome AAA-ATPase subunit, PROTEASOME REGULATOR1 (PTRE1), and protein synthesis initiation factor eIF2 beta (Supplemental Table S7). Notably, DAS increased after prolonged drought stress (Figure 6B), and was especially pronounced for *GLYCINE RICH PROTEIN 7* (*ATGRP7*). Differential splicing of transcripts of the three phytochrome genes *PHYB*, *PHYC* and *PHYE*, together with *PHYTOCHROME INTERACTING FACTOR 3* (*PIF3*), *PIF3-LIKE 5* and *6*, and *FYPP3* (coding for phy-associated protein phosphatase 3) is particularly striking. PhyB acts in multiple environmental stress pathways (Kim et al., 2021), and PhyB positively mediates drought tolerance (Gonzalez et al., 2012). After subtraction of the genes that undergo DAS specifically in *pp7l* (146 genes), 11% of the 16,850 expressed genes were differentially alternatively spliced. In total, 3,283 transcript isoforms were detected that whose splicing patterns were altered in at least one Col-0 contrast group. GO analysis of the corresponding loci revealed enrichments of transcripts for proteins that were localized to the nuclear speck, cytoplasmic vesicle (for example, the AUTOPHAGY group of ubiquitin-like superfamily proteins – ATG8B, E and F, and ATG13) and the spliceosomal complex itself (Figure 6C). In this respect, several coverage plots suggested reduced splicing efficiency especially after 15 days of drought stress (for example Figure 4). Indeed, this is supported by a reduction in the percentage of reads covering the canonical GT-AG junction (Figure 6D).

These results suggest the importance of alternative splicing of nuclear genes in the modulation of responses to drought stress.

**Figure 6.**
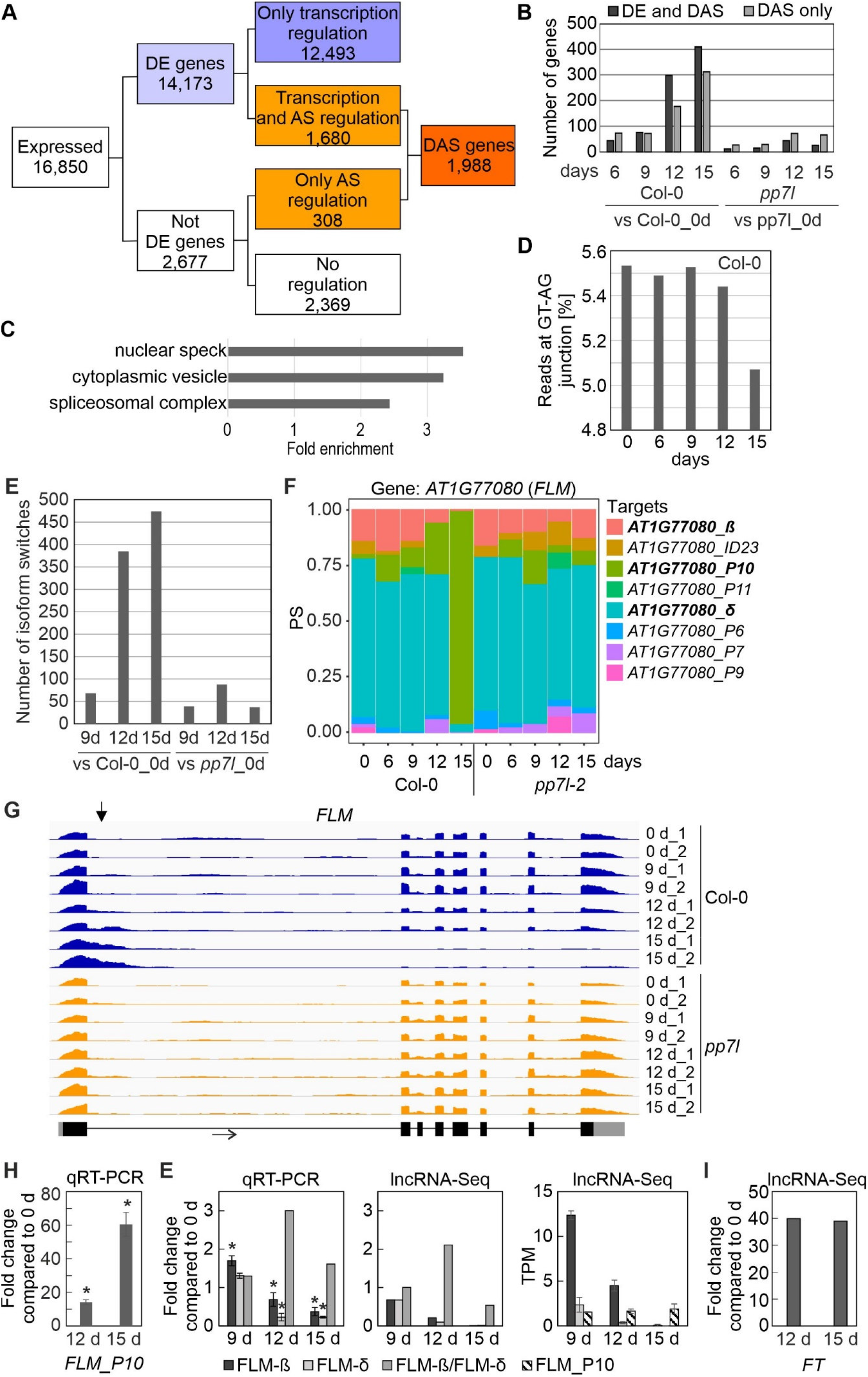
Impact of drought stress on differential alternative splicing (DAS) and transcript isoform switches (ISs). DAS and IS were calculated as described in Materials and Methods. **A,** Flow chart showing the distribution of differentially expressed (DE) and DAS genes. **B,** Number of DE and DAS genes in Col-0 and *pp7l* strains exposed to drought stress compared to the control condition (0 days). **C,** GO enrichment analysis of the cellular component category according to DAVID (Huang da, sherman, 2009). GO terms with a >2-fold change and a Benjamini corrected value of <0.05 are shown. **D,** Percentage of reads spanning the canonical GT-AG splice junction in Col-0 under control (0 days) and drought conditions. **E,** Numbers of ISs detected in Col-0 and *pp7l* after 9, 12 and 15 days of drought stress. **F,** Expression profiles of *FLOWERING LOCUS M* (*FLM*) at the whole-gene level (*AT1G77080*) and at the level of detected transcript isoforms are shown. Arrows mark IS events. PS, percentage of expressed transcripts spliced. **G,** Pattern of *FLM* transcript accumulation. The normalized read depths of transcripts detected in Col-0 and *pp7l* plants under control and prolonged drought conditions were visualized with the Integrative Genomics Viewer (IGV). The vertical arrows indicate differentially spliced regions. Exons (black boxes), introns (black lines) and the 5′- and 3′-UTRs (gray boxes) are shown. **H,** Quantitative reverse transcriptase-polymerase chain reaction (qRT-PCR) analysis of Col-0 control plants (0 d) and plants subjected to drought stress for 12 and 15 days (12 d and 15 d). qRT-PCR was performed with primers specific for the *FLM-β* and FLM_P10 isoforms, and for *AT3G58500* [encoding PROTEIN PHOSPHATASE 2A-4 (PP2A- 4)], which served as a control because it did not significantly change its expression level under drought. Expression values are reported relative to the corresponding transcript levels in non- stressed Col-0. The results were normalized with respect to the expression level of *PP2A-4*. Bars indicate standard deviations. Statistically significant differences (*P* < 0.05) between stressed and non-stressed samples are indicated by an asterisk. Note that the *FLM-β* primers probably detect non-canonical splice forms, in addition to *FLM-β* (Sureshkumar et al., 2016). **I,** Fold changes of *FLM-β* and *FT* transcripts were extracted from lncRNA-Seq results.

### Isoform switch analysis – usage of different isoforms during the course of drought stress

We then sought to identify DAS genes that showed isoform switches (ISs), i.e. in which the relative abundance of different isoforms was altered over the course of drought exposure. In total, 1091 significant (*P* < 0.01) ISs that involved abundant transcript isoforms were detected. Of these, 927 (covering 404 gene loci) were detected in Col-0 when the different time points of drought stress were compared to the control (Figure 6E; Supplemental Table S8). The majority of ISs (more than 450) were identified after 15 days of drought stress. Among the transcripts undergoing ISs were the above-mentioned *LSD1* and *PHYB*. Other examples include *CARBAMOYL PHOSPHATE SYNTHETASE B* (*CARB*, also named *VENOSA 3*), *ZINC-INDUCED FACILITATOR 2* (*ZIF2*), pre-mRNA processing *PRP39A*, and *FLOWERING LOCUS M* (*FLM*) (Figure 6F, Supplemental Figure S6). The ISs involved different types of AS events, which either generated isoforms that encoded different protein variants, or occurred in the 5′ or 3′UTR, without changing the sequence of the protein (Supplemental Table S8). IS events were also detected in transcripts that did not encode a functional protein, e.g., they contained a premature stop codon or did not contain the start codon (Supplemental Table S8). For example, the s3 isoform of the *CARB* transcript that accumulated under prolonged drought stress is predicted not to be translated, as is the P5 isoform of *PRP39A*, and the predicted s3 isoform is very short (Supplemental Figure S7).

The expression behavior of *FLM* transcript isoforms is especially noteworthy. As shown by analysis of lncRNA-Seq data (Figure 6, F and G) and confirmatory qRT-PCR (Figure 6H), the P10 isoform accumulated to very high levels during drought treatment in Col-0 (15-fold and 60-fold after 12 and 15d_DS, respectively, compared to 0d_DS) at the expense of the *FLM-β* and *FLM-δ* isoforms (Figure 6F) which was confirmed by qRT-PCR (Figure 6H). The P10 form covers the first exon and part of the first intron (Figure 6G) resulting in a premature stop codon (Supplemental Table S8). It was suggested that FLM-β is the functional protein that can prevent precocious flowering by repressing the expression of the key flowering time regulator FLOWERING LOCUS T (FT, florigen) (summarized in: (Zarnack et al., 2020; Quiroz et al., 2021). We observed a 40-fold induction of *FT* transcript levels after 12 and 15 days of drought stress compared to 0d_DS (Figure 7I), but after 15 days of drought treatment *FLM-β* and *-δ* expression level were nearly undetectable, suggesting that lack of FLM protein is sufficient to confer the early flowering phenotype under drought conditions (see Figure 7, Discussion section).

**Figure 7.**
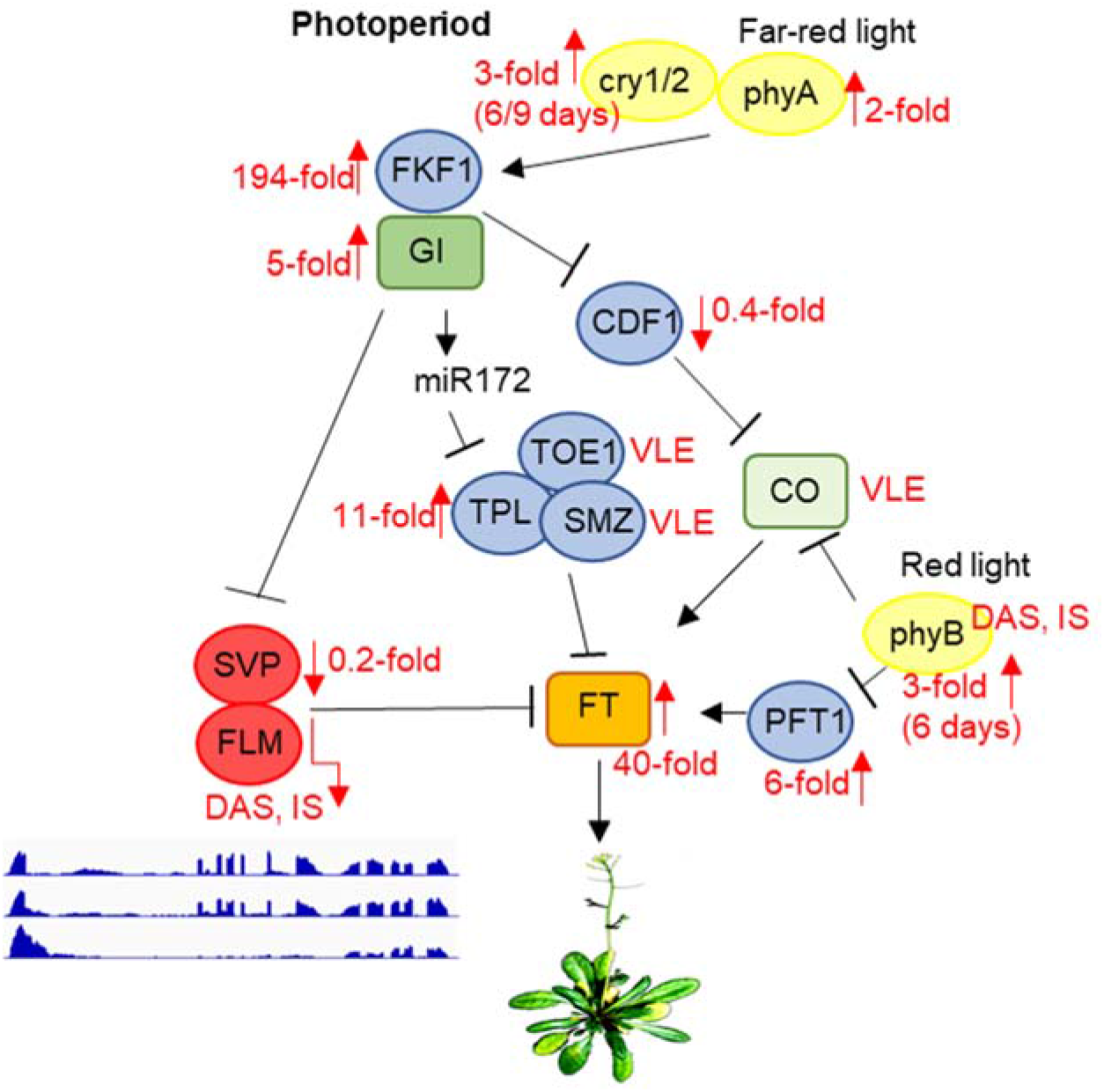
Scheme depicting the photoperiod-dependent flowering pathway, together with inputs from light receptors. Note that the components of the flowering pathway are mainly regulated through transcriptional activation or repression. SVP and FLM are components of the ambient-temperature, GA and autonomous pathways. GI acts in both CO-dependent (by suppressing CDF1) and CO-independent branches by either repressing SVP or by promoting *miR172* expression to control flowering. Flowering itself is ultimately mediated by FT, which is an inducer of flowering. phyB acts to suppress CO protein activity, whereas phyA, cry1, and cry2 function to enhance the activity of CO. In parallel, phyB affects *FT* transcription by suppressing *PFT1*, an upstream activator of *FT*. Black arrows and T-ends indicate positive or negative regulation, respectively. Red arrows together with numbers indicate fold changes after drought stress. *FLM* and *PHYB* transcripts are subject to DAS and IS. Only the main regulatory genes are shown here. The complete flowering time network involves several hundred genes, and is available on the WikiPathways website (https://www.wikipathways.org/index.php/Pathway:WP2312). Modified after Kim (2020) and Leijten et al. (2018). DAS, differential alternative splicing; IS, isoform switching; temp., temperature; VLE, very low expressed.

In conclusion, RNA-Seq analysis uncovered numerous DAS and IS events. In particular, *FLM* isoform switching provides a possible explanation for the early flowering phenotype under drought stress.

## DISCUSSION

To investigate the impact of drought stress on the (post)transcriptome, we included the *pp7l* mutant of which we showed that it can survive long periods of drought stress (see Figure 1). PP7L seems to participate in several stress responses (Xu et al., 2019). PP7L interacts with plant mobile proteins MAINTENANCE OF MERISTEMS (MAIN) and MAIN-LIKE 1 (MAIL1) to silence a subset of transposable elements (de Luxan-Hernandez et al., 2020; Nicolau et al., 2020), but the exact molecular function of those proteins is still unclear. Here we want to emphasize, that it was not our intention to clarify PP7L’s function in this study, but used (post)-transcriptomics behavior of the *pp7l* mutant for comparison with that of the wild-type Col-0. In that way we suggest for example that enhanced editing of chloroplast *rps12* is not a drought-specific effect, because it is enhanced in both Col-0 and *pp7l* (see Figure 5). However, the here generated *pp7l* lncRNA-Seq data might help to dissect the function of PP7L in the future.

As a drought set-up, we used soil-grown plants from which water was withheld for various periods. Many drought models are available, but there is no “ideal” model that can meet all the requirements for drought studies, and individual methods have their particular limitations (Lawlor, 2013). Aqueous or agar media are more stable and reproducible (Ito et al., 2006; Geissler and Wessjohann, 2011; Park et al., 2013) than soil-based models. However, the latter more closely mimic actual drought conditions in the field, which makes it the preferred system in many studies. One typical soil-based model is the “water-withheld-setup”, in which plants are deprived of water until symptoms of wilting are observed (Ali et al., 2020). We deliberately chose this system, because it is accessible to all laboratories. Moreover, field conditions do not allow for the strict control of water availability. To test the robustness of this system, we followed the method used by (Kim et al., 2017). Despite the usage of different methodologies to analyze transcriptomes (lncRNA-Seq vs. microarrays), different control lines (Col-0 versus a Col-0 line containing a *DR5*::*GUS* construct), different sampling (whole plants vs. aerial parts), the transcriptome changes induced by prolonged drought stress detected in the two studies were quite similar (see Fig. 2c).

### Alternative splicing enhances proteome diversity to counteract stress responses

Differential alternative splicing (DAS), which enables multiple transcripts (and therefore proteins with different properties) to be produced from single genes, turns out to be an important aspect of responses to adverse conditions (Laloum et al., 2018) – and components of the spliceosome (which mediates DAS) are known to be altered during drought stress (Marondedze et al., 2019). Here, we identified nearly 2,000 DAS genes (see Figure 6 and Supplemental Table S7), of which 15% were regulated solely at the level of alternative splicing (AS) rather than by alteration of transcription rates, so that these transcripts would have gone unnoticed in a microarray-based approach. Some of the identified genes have already been shown to encode proteins involved in stress pathways. Thus, phyB, LESION SIMULATING DISEASE 1 (LSD1) and GLYCINE RICH PROTEIN 7 (GRP7) have previously been implicated in survival of Arabidopsis plants under drought stress (Gonzalez et al., 2012; Szechynska-Hebda et al., 2016) or higher grain yields of rice under drought conditions (Yang et al., 2014). However, it was not known until now that these effects are mediated by different transcript isoforms. Interestingly, GRP7 itself regulates AS (Streitner et al., 2012). Moreover, CONSTITUTIVELY STRESSED 1 (COST1) regulates autophagy to enhance plant drought tolerance (Bao et al., 2020). In this respect it is remarkable that *ATG8B*, -*E* and -*F*, and *ATG13* (all of which code for AUTOPHAGY ubiquitin-like superfamily proteins) are subject to DAS under drought.

One known example of DAS under drought is that of the ZINC-INDUCED FACILITATOR-LIKE 1 (ZIFL1) transporter. The full-length isoform is localized in the tonoplast of root cells and regulates transport of auxin, but a truncated variant is targeted to the plasma membrane of leaf stomatal guard cells and mediates drought tolerance (Remy et al., 2013). A homolog, *ZINC-INDUCED FACILITATOR 2* (*ZIF2*), is known to produce two splice variants, *ZIF2.1* and *ZIF2.2*, which encode the same proteins, but an intron retention event in the 5’UTR in *ZIF2.2* enhances translation in a zinc-responsive manner and promotes zinc tolerance (Remy et al., 2013). This makes *ZIF2* an attractive candidate – among the many promising DAS genes expressed under drought conditions – for further investigation of the contribution of DAS forms to drought tolerance. Another interesting candidate is *ALDEHYDE DEHYDROGENASE 7B4* (*ALDH7B4*; 3000-fold induced after 12 and 15 days) which also undergoes DAS under drought. Indeed, *aldh7b4* knockout mutants exhibit higher sensitivity to dehydration and salt than do wild-type plants (Kotchoni et al., 2006).

### Organellar (post)transcriptomes under stress

Mitochondria and plastids integrate signals to link metabolic processes with environmental sensing (Dopp et al., 2021). Despite the transfer of most of their genes to the nucleus during evolution, these organelles retain their own gene-expression machinery, thus necessitating tight coordination between organellar and nuclear gene expression (OGE and NGE), which is achieved by retrograde and anterograde signaling (Kleine and Leister, 2016; Dopp et al., 2021). Indeed, under drought, adjustment of organellar and nuclear genomes became apparent at the whole-gene expression level. Drought stress has an overall negative impact on chloroplast transcript accumulation (see Figure 5), which is in line with the down-regulation of nuclear transcripts coding for proteins involved in chloroplast translation and photosynthesis (see cluster 1 in Figure 3). Interestingly, mitochondrial transcripts tended to be slightly upregulated (see Supplemental Figure S5), e.g., those encoding components of succinate-dehydrogenase complex II and enzymes of the tricarboxylic acid cycle (TCA) (see cluster 6 in Figure 3).

Apart from sensing environmental changes, chloroplasts are also targets of adverse conditions (Kleine et al., 2021), and it has been shown that the accumulation of specific plastid RNAs is regulated in mutants with photosynthetic defects and in plants exposed to stresses (see for instance (Cho et al., 2009). Much effort has been put into the investigation of changes in NGE in response to altered organellar states, at both the whole-gene and more recently post- transcriptome levels (Petrillo et al., 2014; Fang et al., 2019; Tiwari et al., 2020). However, to the best of our knowledge, only one study of Arabidopsis organellar post-transcriptomes in chloroplasts has been reported previously (Castandet et al., 2016) and there are none for mitochondria. In the context of the development of a chloroplast RNA-Seq bioinformatics pipeline, an analysis of suitable (i.e. lncRNA-Seq) published transcriptomes (Di et al., 2014) revealed a global reduction in chloroplast splicing and editing efficiency, and an increased abundance of transcripts in response to heat, while short-term “drought” treatment (3 and 12 h in 300 mM mannitol) had negligible effects on the chloroplast (post)transcriptome (Castandet et al., 2016). It should be mentioned that sequencing was performed only in one copy and these data are not statistically sound. Defects in editing or splicing can have profound effects on plant development, and even result in lethality (Kleine and Leister, 2015), and altered adaptability to environmental stresses (Leister et al., 2017; Zhang et al., 2020). Therefore, we wanted to investigate the reverse case: What impact does drought have on the overall accumulation, splicing and editing of organellar transcripts?

Interestingly, levels of mitochondrial and chloroplast RNAs under drought were inversely regulated. While amounts of mitochondrial RNAs tended to rise (see Supplemental Figure S5), drought had a profoundly negative impact on the accumulation of chloroplast transcripts (see Figure 5). The most severely affected transcripts – those encoding a component of the large ribosomal subunit (rpl23) and the D3, F and K subunits of the NAD(P)H dehydrogenase (NDH) complex – were reduced to 3% of their starting amounts. In contrast, transcripts coding for mitochondrial NADH dehydrogenase subunits increased during prolonged drought stress (see Supplemental Figure 6), and the two most repressed mitochondrial transcripts after 15 days of drought stress were *ATP synthase subunit 1* (*atp1*; 0.3%) and *rpsL2* (17.8%). The function of the NDH complex has been extensively discussed and has yet to be resolved (Labs et al., 2016; Yamori and Shikanai, 2016). In particular, the role of the NDH complex under different stress conditions remains controversial (Yamori and Shikanai, 2016; Lenzen et al., 2020).

Editing capacity in Col-0 mitochondria under drought was not changed (see Supplemental Figure S5), but a reduced editing capacity was observed especially for the chloroplast *ndhD*, *ndhF*, *rpl23* transcripts, and *rpoA* and *rpoB* transcripts coding for the α and β subunits of the plastid-encoded RNA polymerase (PEP) (see Figure 5). Note here that *ndhD*, *rpoA* and *rpoB* are not among the most reduced transcripts, implying that editing of these transcripts plays a more prominent role than their accumulation under drought conditions. Furthermore, editing capacity was maintained or even enhanced until the sudden drop occurred after 15 days of drought stress. This explains the maintenance of relatively high levels of *psbA*, -*C* and -*D* transcripts, because non-functional editing of *rpoB* transcripts leads to a significant decrease in PEP activity (Zhou et al., 2009).

In mitochondria, the only detected down-regulation of splicing efficiency was transiently observed for *cytochrome oxidase 2* (*cox2*), *nad1* and *nad7-2*, and this was slightly enhanced after 15 days (see Supplemental Figure S5). In chloroplasts also, splicing efficiency was not generally reduced, but slightly enhanced after 15 days of drought (see Figure 5). Thus, organelle-specific splicing seems to be of marginal importance for the drought stress response.

### Early flowering under drought stress is most probably caused by a lack of functional FLM

The drought escape (DE) strategy involves an earlier switch from vegetative to reproductive development, enabling reproduction before severe water deficit prohibits plant survival (Kenney et al., 2014). Early flowering as a DE mechanism is extremely important and research on the topic has a very long history (Shavrukov et al., 2017). Under a 12 h/12 h light/dark regime, ecotypes with low expression of *FRIGIDA* (*FRI*), or a null *FRI* allele as in Col-0, confer early flowering under drought (Lovell et al., 2013). Also, in our experimental setup in which we used long-day conditions, Col-0 plants began to flower earlier under drought. In Arabidopsis, flowering is ultimately achieved by activating expression of the gene *FLOWERING LOCUS T* (*FT*) (Searle and Coupland, 2004), which in turn activates *SOC1* and other meristem-identity genes (Leijten et al., 2018; Kim, 2020). Accordingly, in our experimental setup, *FT* mRNA levels are strongly elevated under drought treatment (40-fold induction after 12 and 15 days, Figure 7, see Supplemental Table S2). Six major pathways have been implicated in the control of flowering time: the vernalization, age, ambient temperature, photoperiodic, autonomous, and gibberellin (GA) pathways (Leijten et al., 2018). Mutants in the autonomous pathway (late flowering irrespective of the photoperiod) and the GA pathway show a similar DE response to the wild type (Riboni et al., 2013). However, the DE response is dependent on the photoperiod – at least for the Arabidopsis Col-0 and L*er* ecotypes – because short-day-grown plants do not flower earlier under drought stress (Riboni et al., 2013). In addition to FT, the photoperiodic pathway is characterized by two other key components, GIGANTEA (GI) and CONSTANS (CO) (Putterill et al., 1995; Fowler et al., 1999). Complete absence of the DE response was observed in *gi* mutants in both the Col-0 and L*er* backgrounds, but this response does not appear to require the activity of CO (Riboni et al., 2013), which is a transcriptional regulator of *FT* that acts downstream of GI. Correspondingly, *GI* mRNA levels are 5-fold induced and those of *CO* barely detectable under our drought conditions (Figure 7). The precise role of GI action in the DE response has not yet been clarified (Riboni et al., 2013). GI also acts in CO-independent branches by either directly binding to the FT promoter and competing with some repressors of *FT* such as SHORT VEGETATIVE PHASE (SVP), or promoting *miR172* expression which inhibits the expression of AP2-like transcription factors, such as *TOE1*, *TPL*, *SMZ*, thus repressing *FT* transcription (summarized in (Kim, 2020). Under our drought conditions, *TOE1* and *SMZ* mRNAs are barely detectable, while levels of *TPL* are elevated 11-fold (Figure 7). These results do yield a clear picture of the respective roles of these proteins. However, amounts of *SVP* mRNAs are reduced to 20% of their starting levels in accordance with the elevated *GI* and *FT* levels. However, the alternative splicing and isoform switching (IS) of *FLOWERING LOCUS M* (*FLM*) (see Figure 6), another repressor of *FT* transcription which also acts independently of CO (Balasubramanian et al., 2006), is particularly noteworthy. FLM and SVP also participate in the autonomous (Scortecci et al., 2001) and the thermosensory (Balasubramanian et al., 2006) flowering pathways. *FLM* undergoes temperature-dependent alternative splicing, and the roles of the different isoforms have been extensively and controversially discussed in the literature. It was proposed that the major isoforms, *FLM-β* and *FLM-δ*, which result from the alternative usage of exons 2 (*FLM-β*) and 3 (*FLM-δ*) (Lee et al., 2013; Pose et al., 2013) might compete for interaction with SVP, and the SVP-FLM-β complex is predominantly formed at low temperatures and prevents precocious flowering (Pose et al., 2013). One model for flowering at high temperatures suggests that degradation of SVP reduces the abundance of the SVP-FLM-β repressor complex (Lee et al., 2013), the other model is based on the idea that a higher *FLM-δ*/*FLM-β* ratio favors the formation of a SVP–FLM-δ complex that is impaired in DNA binding and acts as a dominant-negative activator of flowering at higher temperatures (Pose et al., 2013). We assume that under (our) drought conditions a higher FLM-*β*/FLM-*δ* ratio is unlikely to be the trigger for the flowering DE response. Our qRT-PCR data suggest that amounts of the *FLM-β* isoform increase after 9 days of drought stress, while lncRNA-Seq analysis indicates that levels of FLM-ß are slightly reduced relative to the onset of drought treatment (see Figure 6). This discrepancy can be explained by accumulation of other non-functional isoforms containing exon 2 or exon 3, respectively (see also Sureshkumar et al. (2016), which are amplified by qRT-PCR owing to the frequent use of primer designs in which the reverse primer is situated in exon 2 (*FLM-ß*) or 3 (*FLM-δ*), respectively (for example also used in Pose et al. (2013)). Although the *FLM-β*/*FLM-δ* ratio rises especially after 12 days of drought stress, the accumulation of these forms is very low in comparison to that prior to drought stress, and both are nearly undetectable anymore after 15 days. Moreover, the *FLM_P10* form, which is predominantly produced under prolonged drought stress, contains a premature stop codon that would encode a protein of only 62 amino acids (see Supplemental Table S8). Indeed, under high temperatures too, *FLM* expression is downregulated by AS coupled with nonsense-mediated mRNA decay (AS-NMD), and the majority of non-canonical *FLM* transcripts contained premature termination codons that would result in truncated proteins of less than 100 amino acids (Sureshkumar et al., 2016). All in all, we suggest that drought-mediated early flowering under long-day conditions is caused by the absence of functional FLM protein and not by the ratio of isoforms.

The decision whether to investigate under long- or short-day conditions is a very important one, since it may result in contrasting outcomes. In relation to drought research, SVP was shown to be up-regulated at the mRNA level and to confer drought tolerance in short-day-grown plants (Bechtold et al., 2016; Wang et al., 2018). However, in our long-day kinetic experiment, SVP is downregulated at all times, and would not have been discovered as a positive factor in drought stress.

Moreover, our transcriptomic data sets provide a rich resource for the elucidation of the many facets of drought stress mechanisms. In particular, evaluation of the relative contributions of different splicing isoforms to drought tolerance will increase our understanding of the modulation of abiotic stress responses, thus enabling the development of new strategies to improve plant performance under adverse environmental conditions.

## METHODS

### Plant material and growth conditions

All Arabidopsis (*Arabidopsis thaliana*) lines used in this study share the Columbia genetic background. Seeds were sown on 1/2 MS medium containing 1% (w/v) sucrose and 0.8% (w/v) agar, incubated at 4°C for 2 d, and transferred to a climate chamber under a 16-h-light/8-h-dark cycle with a light intensity of 100 μE m^-2^ s^-1^ at 22°C. For physiological experiments, the plants were grown on potting soil (A210, Stender, Schermbeck, Germany) for 3 or 4 weeks.

The mutants analyzed in this study included *pp7l-1*, *pp7l-2*, *cry1cry2*, *cop1-4*, which were used previously (Xu et al., 2019). For drought treatments, 7-d-old seedlings were transferred to pots, and grown for a further 2 weeks under normal growth conditions. Three-week-old plants were subjected to drought conditions by withholding water for the indicated times. The drought phenotypes were documented with a camera (Canon 550D, Krefeld, Germany).

### Chlorophyll *a* Fluorescence Measurements

Chlorophyll fluorescence parameters were measured using an imaging Chl fluorometer (Imaging PAM, M-Series; Walz, Effeltrich, Germany). The method employed involved dark adaptation for 30-min, followed by the determination of Fv/Fm. F_0_ was measured at a low frequency of pulse-modulated measuring light (4 Hz, intensity 3, gain 3, damping 2), while Fm was quantified following saturation pulses of approximately 2700 µmol photons m^−2^ s^−1^ for 0.8 s. The calculations and plotting of the parameters were performed using the ImagingWinGigE software.

### Nucleic Acid Extraction

Leaf tissue (100 mg) was homogenized in extraction buffer containing 200 mM Tris/HCl (pH 7.5), 25 mM NaCl, 25 mM EDTA and 0.5% (w/v) SDS. After centrifugation, DNA was precipitated from the supernatant by adding isopropyl alcohol. After washing with 70% (v/v) ethanol, the DNA was dissolved in distilled water.

For RNA isolation, plant material was harvested, frozen in liquid nitrogen and crushed in a TissueLyser (Retsch, model: MM400). Total RNA was extracted with the Direct-zol RNA Kit (Zymo Research, Irvine, USA) according to the manufacturer’s protocol. RNA samples were quantified with a Nanodrop spectrophotometer (ThermoFisher Scientific, Langenselbold, Germany) and RNA integrity and quality were assessed with an Agilent 2100 Bioanalyzer (Agilent, Santa Clara, USA). Isolated RNA was stored at −80 °C until further use.

### cDNA Synthesis and Quantitative Reverse Transcriptase-Polymerase Chain Reaction (qRT-PCR) Analysis

One microgram of RNA was employed to synthesize cDNA using the iScript cDNA Synthesis Kit (Bio-Rad, Munich, Germany). Quantitative real-time PCR analysis was performed on a CFX Connect™ Real-Time PCR Detection System (Bio-Rad) with the SsoAdvanced™ Universal SYBR® Green Supermix (Bio-Rad). The gene-specific primers used for this assay are listed in Supplemental Table 9. Each sample was quantified in triplicate and normalized using *AT3G58500* (which codes for protein phosphatase 2A-4 and showed minimal expression changes during drought treatment) as an internal control.

### Long Non-Coding RNA-Sequencing (lncRNA-Seq)

Total RNA from plants was isolated using TRIzol Reagent™ (Thermo Fisher Scientific, Waltham, MA, USA) and purified using an RNA Clean & Concentrator (Zymo Research, Irvine, USA) according to the manufacturer’s instructions. RNA integrity and quality were assessed with an Agilent 2100 Bioanalyzer (Santa Clara, USA). RNA was depleted of ribosomal RNA with the RiboMinus Plant Kit for RNA-seq (Thermo Fisher Scientific). The purified RNAs were first fragmented randomly to short fragments of 150-200 bp by addition of fragmentation buffer, followed by cDNA synthesis using random hexamers. After the first strand was synthesized, second-strand synthesis buffer (Illumina), dNTPs (dUTP, dATP, dGTP and dCTP) and DNA polymerase I were added to synthesize the second-strand. This was followed by purification by AMPure XP beads, terminal repair, polyadenylation, sequencing adapter ligation, size selection and degradation of second-strand U-contained cDNA by the USER enzyme. 150-bp paired-end sequencing was conducted on an Illumina HiSeq 2500 system (Illumina, San Diego, USA) at Novogene Biotech (Beijing, China) with standard Illumina protocols. Three independent biological replicates were used per genotype (Col-0 and *pp7l*) and condition (0, 6, 9, 12 and 15 days). However, library preparation from one sample (Col-0 subjected to 6 days of drought treatment) failed. In this case, two replicates were used. Sequencing data have been deposited to Gene Expression Omnibus (Edgar et al., 2002).

### Chloro-Seq analysis

To detect instances of editing and splicing of organellar transcripts in our lncRNA-Seq data, a modified Chloro-Seq pipeline (Malbert et al., 2018) was used. The lncRNA-Seq reads were processed on the Galaxy platform (https://usegalaxy.org/) to remove the adaptors by Trimmomatic (Bolger et al., 2014), then sequencing quality was assessed with FastQC (http://www.bioinformatics.babraham.ac.uk/projects/fastqc/). The trimmed reads were mapped using RNA STAR (Dobin et al., 2013) with TAIR10_Chr.all.fasta (https://www.arabidopsis.org/download_files/Genes/TAIR10_genome_release/TAIR10_chromosome_files/TAIR10_chr_all.fas) as reference genome file, and Araport11_GFF3_genes_transposons.201606.gff (https://www.arabidopsis.org/download_files/Genes/Araport11_genome_release/archived/Araport11_GFF3_genes_transposons.201606.gff.gz) as gene model file for splice junctions. The mapped bam file was later used as the input for Chloro-Seq analysis (https://github.com/BenoitCastandet/chloroseq) to determine editing and splicing efficiency.

### 3D RNA-Seq analysis

To quantify overall transcript accumulation and detect alternative splicing (AS) of nuclear transcripts, the 3D RNA-Seq pipeline (Guo et al., 2021) was used. To this end, RNA-Seq reads were prepared on the Galaxy platform (https://usegalaxy.org/). Adaptors were removed with Trimmomatic (Bolger et al., 2014), and sequencing quality was accessed with FastQC (http://www.bioinformatics.babraham.ac.uk/projects/fastqc/). Transcript abundances were calculated using Salmon (Patro et al., 2017) and AtRTD2-QUASI as reference transcriptome (Zhang et al., 2017). The generated files were uploaded into the 3D RNA-Seq app (https://3drnaseq.hutton.ac.uk/app_direct/3DRNAseq; (Calixto et al., 2018; Guo et al., 2021) and transcript per million reads (TPMs) were calculated using the implemented lengthScaledTPM method. Weakly expressed transcripts and genes were filtered out based on the mean-variance trend of the data. The expected decreasing trend between data mean and variance was observed when expressed transcripts were determined as ≥ 1 of the 29 samples with count per million reads (CPM) ≥ 4, which provided an optimal filter of low expression. A gene was declared to be expressed if any of its transcripts met the above criteria. The TMM method was used to normalize the gene and transcript read counts to log_2_-CPM, and a principal component analysis (PCA) plot showed that the RNA-seq data did not have distinct batch effects. The log_2_ fold change (FC) of gene/transcript abundances was calculated based on contrast groups and the significance of expression changes was determined using t-test. A gene/transcript was significantly differentially expressed (DE) in a contrast group if it had an absolute log_2_ FC ≥ 1 and an adjusted *P*-value < 0.01 after applying multiple testing (Benjamini et al., 2001). At the alternative splicing level, DTU (differential transcript usage) transcripts were identified by comparing the log_2_ FC of a transcript to the weighted average of log_2_ FCs (weights were based on their standard deviation) of all remaining transcripts of the same gene. A transcript was declared to exhibit significant DTU if it had an adjusted *P*-value < 0.01 and a Δ Percent Spliced (ΔPS) ratio ≥ 0.1. For DAS genes, each individual transcript log_2_ FC was compared to the gene-level log_2_ FC, which was calculated as the weighted average of log_2_ FCs of all transcripts of the gene. Then P-values of individual transcript comparison were summarized to a single gene-level *P*-value with an F-test. A gene was significantly DAS in a contrast group if it had an adjusted *P*-value < 0.01 and any of its transcripts had a ΔPS ratio ≥ 0.1.

Transcript isoform switches (ISs) were recognized as such if the order of relative expression levels of a pair of alternatively spliced isoforms underwent a reversal. The Pair- Wise Isoform Switch (isokTSP) method was used to detect the isoform switch points between conditions of contrast groups (except for the 6-days samples). The method defined the ISs between any pair of transcripts within genes using mean values of conditions. It described the significant ISs using five different features of metrics: 1) the probability of switching (i.e. the frequency of samples reversing their relative abundances at the switches) was set to > 0.5; (2) the sum of the average differences of the two isoforms in both intervals before and after the switch point was set to ΔTPM > 1; (3) the significance of the differences between the switched isoform abundances before and after the switch was set to a BH-adjusted *P*-value < 0.01; (4) both of the interval lengths before and after switch were set to 1; and (5) the Pearson correlation of two isoforms was set to >0 (see Guo et al. (2021) for more details).

### Detection of accumulation of transposable elements

Reads were mapped to the Arabidopsis genome (TAIR10) with the gapped-read mapper RNA Star (Galaxy Version 2.7.8a+galaxy0; (Dobin et al., 2013). Reads covering regions of transposable elements were counted with featureCounts (Liao et al., 2014) with the help of the GFF3 gene annotation in Araport11 (www.araport.org/data/araport11). Differentially expressed transposable elements were identified with DESeq2 (Love et al., 2014), running with the fit type set to “parametric”, and applying a 2-fold change cut-off and an adjusted *P*-value of < 0.05.

### Visualization of gene expression data

Normalized read depths of transcripts were visualized with the Integrative Genomics Viewer (IGV; (Robinson et al., 2011). The heatmaps presented in Figures 3, 4 and 6 were generated with the help of ClustVis, a web tool for visualizing clustering of multivariate data (Metsalu and Vilo, 2015).

### Determination of Gene Ontology (GO) enrichments

GO enrichments were obtained from the Database for Annotation, Visualization and Integrated Discovery (DAVID; (Huang da et al., 2009), applying a cut-off of 2-fold enrichment compared to the expected frequency in the Arabidopsis genome and an FDR (Benjamini-Hochberg) ≤ 0.05. Non-redundant GO terms were selected in the interface REVIGO using the medium similarity (0.7) parameter (Supek et al., 2011).

## FUNDING

This work was supported by the Deutsche Forschungsgemeinschaft (TRR175, Project C01 to T.K. and Project C05 to D.L.).

## AUTHOR CONTRIBUTIONS

Conceptualization, T.K. and D.X.; Methodology, D.X., Q.T., and T.K.; Validation, D.X. and Q.T.; Formal Analysis, T.K.; Investigation, D.X., Q.T., and T.K.; Resources, D.L. and T.K.; Writing – Original Draft, T.K.; Writing – Review & Editing, Q.T., D.X., D.L., and T.K.; Funding Acquisition, D.L. and T.K.; Supervision, T.K.

## ACKNOWLEDGEMENTS

We thank Ramona Kandler for excellent technical assistance and Paul Hardy for critical reading of the manuscript.

## Conflict of interest statement

The authors declare no conflict of interest.

## SUPPLEMENTAL DATA

Supplemental Figures 1 to S6

Supplemental Tables S1 to S9

### Legends to Supplemental Tables

**Supplemental Table S1.** Sequencing depth of lncRNA-Seq experiments. Long non-coding RNA sequencing (lncRNA-Seq) was performed with RNA isolated from 3-week-old Col-0 (Col) and *pp7l* plants grown under optimal conditions (time-point 0 days; 0d) and after water had been withheld for 6, 9, 12 and 15 days, respectively (6d, 9d, 12d and 15d). Sequences were mapped and analyzed as described in Materials and Methods.

**Supplemental Table S2.** Genes differentially expressed under drought stress. Long non-coding RNA sequencing (lncRNA-Seq) was performed with RNA isolated from 3-week-old Col-0 (Col) and *pp7l* plants grown under optimal conditions (time-point 0 days; 0d) and after water had been withheld for 6, 9, 12 and 15 days, respectively (6d, 9d, 12d and 15d). Sequences were mapped and analyzed with the 3D RNA-Seq pipeline as described in Materials and Methods.

**Supplemental Table S3.** Drought-stress marker genes. Lists of genes whose transcripts are regulated in the same direction (up or down) after 9, 12 and 15 days of drought stress in our dataset as well as in Kim et al. (2017).

**Supplemental Table S4.** Gene Ontology (GO) analysis of the nine clusters listed in Figure 5. Biological process and cellular component categories according to DAVID (Huang da et al., 2009) and REVIGO (Supek et al., 2011).

**Supplemental Table S5.** Normalized read counts across regions containing transposable elements (TEs).

Supplemental Table S6. Abundance of chloroplast and mitochondrial transcripts. (A) Log_2_ fold changes of RNAs encoded by the chloroplast genome. Note that analysis of tRNAs was excluded. (B) Log_2_ fold changes of RNAs encoded by the mitochondrial genome.

**Supplemental Table S7.** List of differentially alternatively spliced (DAS) genes under drought stress.

**Supplemental Table S8.** List of genes that showed isoform switches (ISs) during drought stress.

**Supplemental Table S9**. Primers used in this study.

## Notes

### Competing Interest Statement

The authors have declared no competing interest.

## REFERENCES

1. Ali S, Hayat K, Iqbal A, Xie L (2020) Implications of Abscisic Acid in the Drought Stress Tolerance of Plants. Agronomy 10: 1323

2. Balasubramanian S, Sureshkumar S, Lempe J, Weigel D (2006) Potent induction of Arabidopsis thaliana flowering by elevated growth temperature. PLoS Genet 2: e106

3. Bao Y, Song WM, Wang P, Yu X, Li B, Jiang C, Shiu SH, Zhang H, Bassham DC (2020) COST1 regulates autophagy to control plant drought tolerance. Proc Natl Acad Sci U S A 117: 7482–7493

4. Bashir K, Matsui A, Rasheed S, Seki M (2019) Recent advances in the characterization of plant transcriptomes in response to drought, salinity, heat, and cold stress. F1000Res 8

5. Bechtold U, Penfold CA, Jenkins DJ, Legaie R, Moore JD, Lawson T, Matthews JS, Vialet-Chabrand SR, Baxter L, Subramaniam S, Hickman R, Florance H, Sambles C, Salmon DL, Feil R, Bowden L, Hill C, Baker NR, Lunn JE, Finkenstadt B, Mead A, Buchanan-Wollaston V, Beynon J, Rand DA, Wild DL, Denby KJ, Ott S, Smirnoff N, Mullineaux PM (2016) Time-Series Transcriptomics Reveals That AGAMOUS-LIKE22 Affects Primary Metabolism and Developmental Processes in Drought-Stressed Arabidopsis. Plant Cell 28: 345–366

6. Benjamini Y, Drai D, Elmer G, Kafkafi N, Golani I (2001) Controlling the false discovery rate in behavior genetics research. Behav Brain Res 125: 279–284

7. Bernacki MJ, Rusaczonek A, Czarnocka W, Karpinski S (2021) Salicylic Acid Accumulation Controlled by LSD1 Is Essential in Triggering Cell Death in Response to Abiotic Stress. Cells 10

8. Blum A (2016) Stress, strain, signaling, and adaptation--not just a matter of definition. J Exp Bot 67: 562–565

9. Blum A, Tuberosa R (2018) Dehydration survival of crop plants and its measurement. J Exp Bot 69: 975–981

10. Bolger AM, Lohse M, Usadel B (2014) Trimmomatic: a flexible trimmer for Illumina sequence data. Bioinformatics 30: 2114–2120

11. Calixto CPG, Guo W, James AB, Tzioutziou NA, Entizne JC, Panter PE, Knight H, Nimmo HG, Zhang R, Brown JWS (2018) Rapid and Dynamic Alternative Splicing Impacts the Arabidopsis Cold Response Transcriptome. Plant Cell 30: 1424–1444

12. Castandet B, Hotto AM, Strickler SR, Stern DB (2016) ChloroSeq, an Optimized Chloroplast RNA-Seq Bioinformatic Pipeline, Reveals Remodeling of the Organellar Transcriptome Under Heat Stress. G3 (Bethesda) 6: 2817–2827

13. Cho WK, Geimer S, Meurer J (2009) Cluster analysis and comparison of various chloroplast transcriptomes and genes in Arabidopsis thaliana. DNA Res 16: 31–44

14. de Luxan-Hernandez C, Lohmann J, Hellmeyer W, Seanpong S, Woltje K, Magyar Z, Pettko-Szandtner A, Pelissier T, De Jaeger G, Hoth S, Mathieu O, Weingartner M (2020) PP7L is essential for MAIL1-mediated transposable element silencing and primary root growth. Plant J 102: 703–717

15. Di C, Yuan J, Wu Y, Li J, Lin H, Hu L, Zhang T, Qi Y, Gerstein MB, Guo Y, Lu ZJ (2014) Characterization of stress-responsive lncRNAs in Arabidopsis thaliana by integrating expression, epigenetic and structural features. Plant J 80: 848–861

16. Dobin A, Davis CA, Schlesinger F, Drenkow J, Zaleski C, Jha S, Batut P, Chaisson M, Gingeras TR (2013) STAR: ultrafast universal RNA-seq aligner. Bioinformatics 29: 15–21

17. Dopp IJ, Yang X, Mackenzie SA (2021) A new take on organelle-mediated stress sensing in plants. New Phytol 230: 2148–2153

18. Edgar R, Domrachev M, Lash AE (2002) Gene Expression Omnibus: NCBI gene expression and hybridization array data repository. Nucleic Acids Res 30: 207–210

19. Estavillo GM, Crisp PA, Pornsiriwong W, Wirtz M, Collinge D, Carrie C, Giraud E, Whelan J, David P, Javot H, Brearley C, Hell R, Marin E, Pogson BJ (2011) Evidence for a SAL1-PAP chloroplast retrograde pathway that functions in drought and high light signaling in Arabidopsis. Plant Cell 23: 3992–4012

20. Fang X, Zhao G, Zhang S, Li Y, Gu H, Li Y, Zhao Q, Qi Y (2019) Chloroplast-to-Nucleus Signaling Regulates MicroRNA Biogenesis in Arabidopsis. Dev Cell 48: 371–382 e374

21. Fowler S, Lee K, Onouchi H, Samach A, Richardson K, Morris B, Coupland G, Putterill J (1999) GIGANTEA: a circadian clock-controlled gene that regulates photoperiodic flowering in Arabidopsis and encodes a protein with several possible membrane-spanning domains. EMBO J 18: 4679–4688

22. Geissler T, Wessjohann LA (2011) A Whole-Plant Microtiter Plate Assay for Drought Stress Tolerance-Inducing Effects. Journal of Plant Growth Regulation 30: 504–511

23. Germain A, Hotto AM, Barkan A, Stern DB (2013) RNA processing and decay in plastids. Wiley Interdiscip Rev RNA 4: 295–316

24. Gonzalez CV, Ibarra SE, Piccoli PN, Botto JF, Boccalandro HE (2012) Phytochrome B increases drought tolerance by enhancing ABA sensitivity in Arabidopsis thaliana. Plant Cell Environ 35: 1958–1968

25. Guo W, Tzioutziou NA, Stephen G, Milne I, Calixto CP, Waugh R, Brown JWS, Zhang R (2021) 3D RNA-seq: a powerful and flexible tool for rapid and accurate differential expression and alternative splicing analysis of RNA-seq data for biologists. RNA Biol 18: 1574–1587

26. Gupta A, Rico-Medina A, Cano-Delgado AI (2020) The physiology of plant responses to drought. Science 368: 266–269

27. Hong Y, Wang Z, Liu X, Yao J, Kong X, Shi H, Zhu JK (2020) Two Chloroplast Proteins Negatively Regulate Plant Drought Resistance Through Separate Pathways. Plant Physiol 182: 1007–1021

28. Hu H, Xiong L (2014) Genetic engineering and breeding of drought-resistant crops. Annu Rev Plant Biol 65: 715–741

29. Huang da W, Sherman BT, Lempicki RA (2009) Bioinformatics enrichment tools: paths toward the comprehensive functional analysis of large gene lists. Nucleic Acids Res 37: 1–13

30. Ito K, Matsukawa K, Kato Y (2006) Functional analysis of skunk cabbage SfUCPB, a unique uncoupling protein lacking the fifth transmembrane domain, in yeast cells. Biochem Biophys Res Commun 349: 383–390

31. Kenney AM, McKay JK, Richards JH, Juenger TE (2014) Direct and indirect selection on flowering time, water-use efficiency (WUE, delta (13)C), and WUE plasticity to drought in Arabidopsis thaliana. Ecol Evol 4: 4505–4521

32. Kim DH (2020) Current understanding of flowering pathways in plants: focusing on the vernalization pathway in Arabidopsis and several vegetable crop plants. Horticulture Environment and Biotechnology 61: 209–227

33. Kim JM, To TK, Matsui A, Tanoi K, Kobayashi NI, Matsuda F, Habu Y, Ogawa D, Sakamoto T, Matsunaga S, Bashir K, Rasheed S, Ando M, Takeda H, Kawaura K, Kusano M, Fukushima A, Endo TA, Kuromori T, Ishida J, Morosawa T, Tanaka M, Torii C, Takebayashi Y, Sakakibara H, Ogihara Y, Saito K, Shinozaki K, Devoto A, Seki M (2017) Acetate-mediated novel survival strategy against drought in plants. Nat Plants 3: 17097

34. Kim JY, Lee JH, Park CM (2021) A Multifaceted Action of Phytochrome B in Plant Environmental Adaptation. Front Plant Sci 12: 659712

35. Kleine T, Leister D (2015) Emerging functions of mammalian and plant mTERFs. Biochim Biophys Acta 1847: 786–797

36. Kleine T, Leister D (2016) Retrograde signaling: Organelles go networking. Biochim Biophys Acta 1857: 1313–1325

37. Kleine T, Maier UG, Leister D (2009) DNA transfer from organelles to the nucleus: the idiosyncratic genetics of endosymbiosis. Annu Rev Plant Biol 60: 115–138

38. Kleine T, Nagele T, Neuhaus HE, Schmitz-Linneweber C, Fernie AR, Geigenberger P, Grimm B, Kaufmann K, Klipp E, Meurer J, Mohlmann T, Muhlhaus T, Naranjo B, Nickelsen J, Richter A, Ruwe H, Schroda M, Schwenkert S, Trentmann O, Willmund F, Zoschke R, Leister D (2021) Acclimation in plants - the Green Hub consortium. Plant J 106: 23–40

39. Kooyers N (2019) Are drought resistance strategies associated with life history strategy? A commentary on: ‘Arabidopsis species deploy distinct strategies to cope with drought stress’. Ann Bot 124: vi–viii

40. Kotchoni SO, Kuhns C, Ditzer A, Kirch HH, Bartels D (2006) Over-expression of different aldehyde dehydrogenase genes in Arabidopsis thaliana confers tolerance to abiotic stress and protects plants against lipid peroxidation and oxidative stress. Plant Cell Environ 29: 1033–1048

41. Labs M, Ruhle T, Leister D (2016) The antimycin A-sensitive pathway of cyclic electron flow: from 1963 to 2015. Photosynth Res 129: 231–238

42. Laloum T, Martin G, Duque P (2018) Alternative Splicing Control of Abiotic Stress Responses. Trends Plant Sci 23: 140–150

43. Lawlor DW (2013) Genetic engineering to improve plant performance under drought: physiological evaluation of achievements, limitations, and possibilities. J Exp Bot 64: 83–108

44. Lee JH, Ryu HS, Chung KS, Pose D, Kim S, Schmid M, Ahn JH (2013) Regulation of temperature-responsive flowering by MADS-box transcription factor repressors. Science 342: 628–632

45. Leijten W, Koes R, Roobeek I, Frugis G (2018) Translating Flowering Time From Arabidopsis thaliana to Brassicaceae and Asteraceae Crop Species. Plants (Basel) 7

46. Leister D, Wang L, Kleine T (2017) Organellar Gene Expression and Acclimation of Plants to Environmental Stress. Front Plant Sci 8: 387

47. Lenzen B, Ruhle T, Lehniger MK, Okuzaki A, Labs M, Muino JM, Ohler U, Leister D, Schmitz-Linneweber C (2020) The Chloroplast RNA Binding Protein CP31A Has a Preference for mRNAs Encoding the Subunits of the Chloroplast NAD(P)H Dehydrogenase Complex and Is Required for Their Accumulation. Int J Mol Sci 21

48. Liao Y, Smyth GK, Shi W (2014) featureCounts: an efficient general purpose program for assigning sequence reads to genomic features. Bioinformatics 30: 923–930

49. Liu J, He Z (2020) Small DNA Methylation, Big Player in Plant Abiotic Stress Responses and Memory. Front Plant Sci 11: 595603

50. Love MI, Huber W, Anders S (2014) Moderated estimation of fold change and dispersion for RNA-seq data with DESeq2. Genome Biol 15: 550

51. Lovell JT, Juenger TE, Michaels SD, Lasky JR, Platt A, Richards JH, Yu X, Easlon HM, Sen S, McKay JK (2013) Pleiotropy of FRIGIDA enhances the potential for multivariate adaptation. Proc Biol Sci 280: 20131043

52. Malbert B, Rigaill G, Brunaud V, Lurin C, Delannoy E (2018) Bioinformatic Analysis of Chloroplast Gene Expression and RNA Posttranscriptional Maturations Using RNA Sequencing. Methods Mol Biol 1829: 279–294

53. Mao J, Zhang YC, Sang Y, Li QH, Yang HQ (2005) From The Cover: A role for Arabidopsis cryptochromes and COP1 in the regulation of stomatal opening. Proc Natl Acad Sci U S A 102: 12270–12275

54. Marondedze C, Thomas L, Lilley KS, Gehring C (2019) Drought Stress Causes Specific Changes to the Spliceosome and Stress Granule Components. Front Mol Biosci 6: 163

55. Metsalu T, Vilo J (2015) ClustVis: a web tool for visualizing clustering of multivariate data using Principal Component Analysis and heatmap. Nucleic Acids Res 43: W566–570

56. Moazzam-Jazi M, Ghasemi S, Seyedi SM, Niknam V (2018) COP1 plays a prominent role in drought stress tolerance in Arabidopsis and Pea. Plant Physiol Biochem 130: 678–691

57. Nicolau M, Picault N, Descombin J, Jami-Alahmadi Y, Feng S, Bucher E, Jacobsen SE, Deragon JM, Wohlschlegel J, Moissiard G (2020) The plant mobile domain proteins MAIN and MAIL1 interact with the phosphatase PP7L to regulate gene expression and silence transposable elements in Arabidopsis thaliana. PLoS Genet 16: e1008324

58. Park HJ, Lee SS, You YN, Yoon DH, Kim BG, Ahn JC, Cho HS (2013) A Rice Immunophilin Gene, OsFKBP16-3, Confers Tolerance to Environmental Stress in Arabidopsis and Rice. Int J Mol Sci 14: 5899–5919

59. Patro R, Duggal G, Love MI, Irizarry RA, Kingsford C (2017) Salmon provides fast and bias-aware quantification of transcript expression. Nat Methods 14: 417–419

60. Petrillo E, Godoy Herz MA, Fuchs A, Reifer D, Fuller J, Yanovsky MJ, Simpson C, Brown JW, Barta A, Kalyna M, Kornblihtt AR (2014) A chloroplast retrograde signal regulates nuclear alternative splicing. Science 344: 427–430

61. Pires MV, Pereira Junior AA, Medeiros DB, Daloso DM, Pham PA, Barros KA, Engqvist MK, Florian A, Krahnert I, Maurino VG, Araujo WL, Fernie AR (2016) The influence of alternative pathways of respiration that utilize branched-chain amino acids following water shortage in Arabidopsis. Plant Cell Environ 39: 1304–1319

62. Popova OV, Dinh HQ, Aufsatz W, Jonak C (2013) The RdDM pathway is required for basal heat tolerance in Arabidopsis. Mol Plant 6: 396–410

63. Pose D, Verhage L, Ott F, Yant L, Mathieu J, Angenent GC, Immink RG, Schmid M (2013) Temperature-dependent regulation of flowering by antagonistic FLM variants. Nature 503: 414–417

64. Putterill J, Robson F, Lee K, Simon R, Coupland G (1995) The CONSTANS gene of Arabidopsis promotes flowering and encodes a protein showing similarities to zinc finger transcription factors. Cell 80: 847–857

65. Quiroz S, Yustis JC, Chavez-Hernandez EC, Martinez T, Sanchez MP, Garay-Arroyo A, Alvarez-Buylla ER, Garcia-Ponce B (2021) Beyond the Genetic Pathways, Flowering Regulation Complexity in Arabidopsis thaliana. Int J Mol Sci 22

66. Remy E, Cabrito TR, Baster P, Batista RA, Teixeira MC, Friml J, Sa-Correia I, Duque P (2013) A major facilitator superfamily transporter plays a dual role in polar auxin transport and drought stress tolerance in Arabidopsis. Plant Cell 25: 901–926

67. Riboni M, Galbiati M, Tonelli C, Conti L (2013) GIGANTEA enables drought escape response via abscisic acid-dependent activation of the florigens and SUPPRESSOR OF OVEREXPRESSION OF CONSTANS. Plant Physiol 162: 1706–1719

68. Robinson JT, Thorvaldsdottir H, Winckler W, Guttman M, Lander ES, Getz G, Mesirov JP (2011) Integrative genomics viewer. Nat Biotechnol 29: 24–26

69. Rorbach J, Bobrowicz A, Pearce S, Minczuk M (2014) Polyadenylation in bacteria and organelles. Methods Mol Biol 1125: 211–227

70. Scortecci KC, Michaels SD, Amasino RM (2001) Identification of a MADS-box gene, FLOWERING LOCUS M, that represses flowering. Plant J 26: 229–236

71. Searle I, Coupland G (2004) Induction of flowering by seasonal changes in photoperiod. EMBO J 23: 1217–1222

72. Shavrukov Y, Kurishbayev A, Jatayev S, Shvidchenko V, Zotova L, Koekemoer F, de Groot S, Soole K, Langridge P (2017) Early Flowering as a Drought Escape Mechanism in Plants: How Can It Aid Wheat Production? Front Plant Sci 8: 1950

73. Singh B, Kukreja S, Goutam U (2018) Milestones achieved in response to drought stress through reverse genetic approaches. F1000Res 7: 1311

74. Singh J, Mishra V, Wang F, Huang HY, Pikaard CS (2019) Reaction Mechanisms of Pol IV, RDR2, and DCL3 Drive RNA Channeling in the siRNA-Directed DNA Methylation Pathway. Mol Cell 75: 576–589 e575

75. Streitner C, Koster T, Simpson CG, Shaw P, Danisman S, Brown JW, Staiger D (2012) An hnRNP-like RNA-binding protein affects alternative splicing by in vivo interaction with transcripts in Arabidopsis thaliana. Nucleic Acids Res 40: 11240–11255

76. Supek F, Bosnjak M, Skunca N, Smuc T (2011) REVIGO summarizes and visualizes long lists of gene ontology terms. PLoS One 6: e21800

77. Sureshkumar S, Dent C, Seleznev A, Tasset C, Balasubramanian S (2016) Nonsense- mediated mRNA decay modulates FLM-dependent thermosensory flowering response in Arabidopsis. Nat Plants 2: 16055

78. Szechynska-Hebda M, Czarnocka W, Hebda M, Bernacki MJ, Karpinski S (2016) PAD4, LSD1 and EDS1 regulate drought tolerance, plant biomass production, and cell wall properties. Plant Cell Rep 35: 527–539

79. Tiwari B, Habermann K, Arif MA, Weil HL, Garcia-Molina A, Kleine T, Muhlhaus T, Frank W (2020) Identification of small RNAs during cold acclimation in Arabidopsis thaliana. BMC Plant Biol 20: 298

80. Wang Q, Liu P, Jing H, Zhou XF, Zhao B, Li Y, Jin JB (2021) JMJ27-mediated histone H3K9 demethylation positively regulates drought-stress responses in Arabidopsis. New Phytol 232: 221–236

81. Wang R, Zhao J, Jia M, Xu N, Liang S, Shao J, Qi Y, Liu X, An L, Yu F (2018) Balance between Cytosolic and Chloroplast Translation Affects Leaf Variegation. Plant Physiol 176: 804–818

82. Wang Z, Wang FX, Hong YC, Yao JJ, Ren ZZ, Shi HZ, Zhu JK (2018) The Flowering Repressor SVP Confers Drought Resistance in Arabidopsis by Regulating Abscisic Acid Catabolism. Molecular Plant 11: 1184–1197

83. Wendte JM, Pikaard CS (2017) The RNAs of RNA-directed DNA methylation. Biochim Biophys Acta Gene Regul Mech 1860: 140–148

84. Wilson PB, Estavillo GM, Field KJ, Pornsiriwong W, Carroll AJ, Howell KA, Woo NS, Lake JA, Smith SM, Harvey Millar A, von Caemmerer S, Pogson BJ (2009) The nucleotidase/phosphatase SAL1 is a negative regulator of drought tolerance in Arabidopsis. Plant J 58: 299–317

85. Xu D, Leister D, Kleine T (2020) VENOSA4, a Human dNTPase SAMHD1 Homolog, Contributes to Chloroplast Development and Abiotic Stress Tolerance. Plant Physiol 182: 721–729

86. Xu D, Marino G, Klingl A, Enderle B, Monte E, Kurth J, Hiltbrunner A, Leister D, Kleine T (2019) Extrachloroplastic PP7L Functions in Chloroplast Development and Abiotic Stress Tolerance. Plant Physiol 180: 323–341

87. Yamori W, Shikanai T (2016) Physiological Functions of Cyclic Electron Transport Around Photosystem I in Sustaining Photosynthesis and Plant Growth. Annu Rev Plant Biol 67: 81–106

88. Yang DH, Kwak KJ, Kim MK, Park SJ, Yang KY, Kang H (2014) Expression of Arabidopsis glycine-rich RNA-binding protein AtGRP2 or AtGRP7 improves grain yield of rice (Oryza sativa) under drought stress conditions. Plant Sci 214: 106–112

89. Zarnack K, Balasubramanian S, Gantier MP, Kunetsky V, Kracht M, Schmitz ML, Strasser K (2020) Dynamic mRNP Remodeling in Response to Internal and External Stimuli. Biomolecules 10

90. Zhang H, Lang Z, Zhu JK (2018) Dynamics and function of DNA methylation in plants. Nat Rev Mol Cell Biol 19: 489–506

91. Zhang R, Calixto CPG, Marquez Y, Venhuizen P, Tzioutziou NA, Guo W, Spensley M, Entizne JC, Lewandowska D, Ten Have S, Frei Dit Frey N, Hirt H, James AB, Nimmo HG, Barta A, Kalyna M, Brown JWS (2017) A high quality Arabidopsis transcriptome for accurate transcript-level analysis of alternative splicing. Nucleic Acids Res 45: 5061–5073

92. Zhang Y, Zhang A, Li X, Lu C (2020) The Role of Chloroplast Gene Expression in Plant Responses to Environmental Stress. Int J Mol Sci 21

93. Zhao C, Liu B, Piao S, Wang X, Lobell DB, Huang Y, Huang M, Yao Y, Bassu S, Ciais P, Durand JL, Elliott J, Ewert F, Janssens IA, Li T, Lin E, Liu Q, Martre P, Muller C, Peng S, Penuelas J, Ruane AC, Wallach D, Wang T, Wu D, Liu Z, Zhu Y, Zhu Z, Asseng S (2017) Temperature increase reduces global yields of major crops in four independent estimates. Proc Natl Acad Sci U S A 114: 9326–9331

94. Zhou W, Cheng Y, Yap A, Chateigner-Boutin AL, Delannoy E, Hammani K, Small I, Huang J (2009) The Arabidopsis gene YS1 encoding a DYW protein is required for editing of rpoB transcripts and the rapid development of chloroplasts during early growth. Plant J 58: 82–96

95. Zhu JK (2002) Salt and drought stress signal transduction in plants. Annu Rev Plant Biol 53: 247–273

