## Supplemental Figure for "Response of the organellar and nuclear (post)transcriptomes of Arabidopsis to drought stress"

**SUPPLEMENTAL DATA**

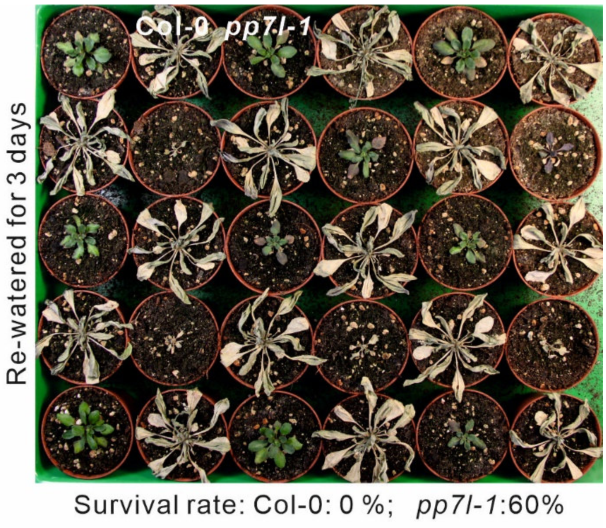


**Supplemental Figure S1.** The *pp7l* mutant can survive long periods of drought stress. Water was with-held for 15 days from 3-week-old Col-0 and *pp7l* mutant plants. Subsequently, plants were re-watered for three days.

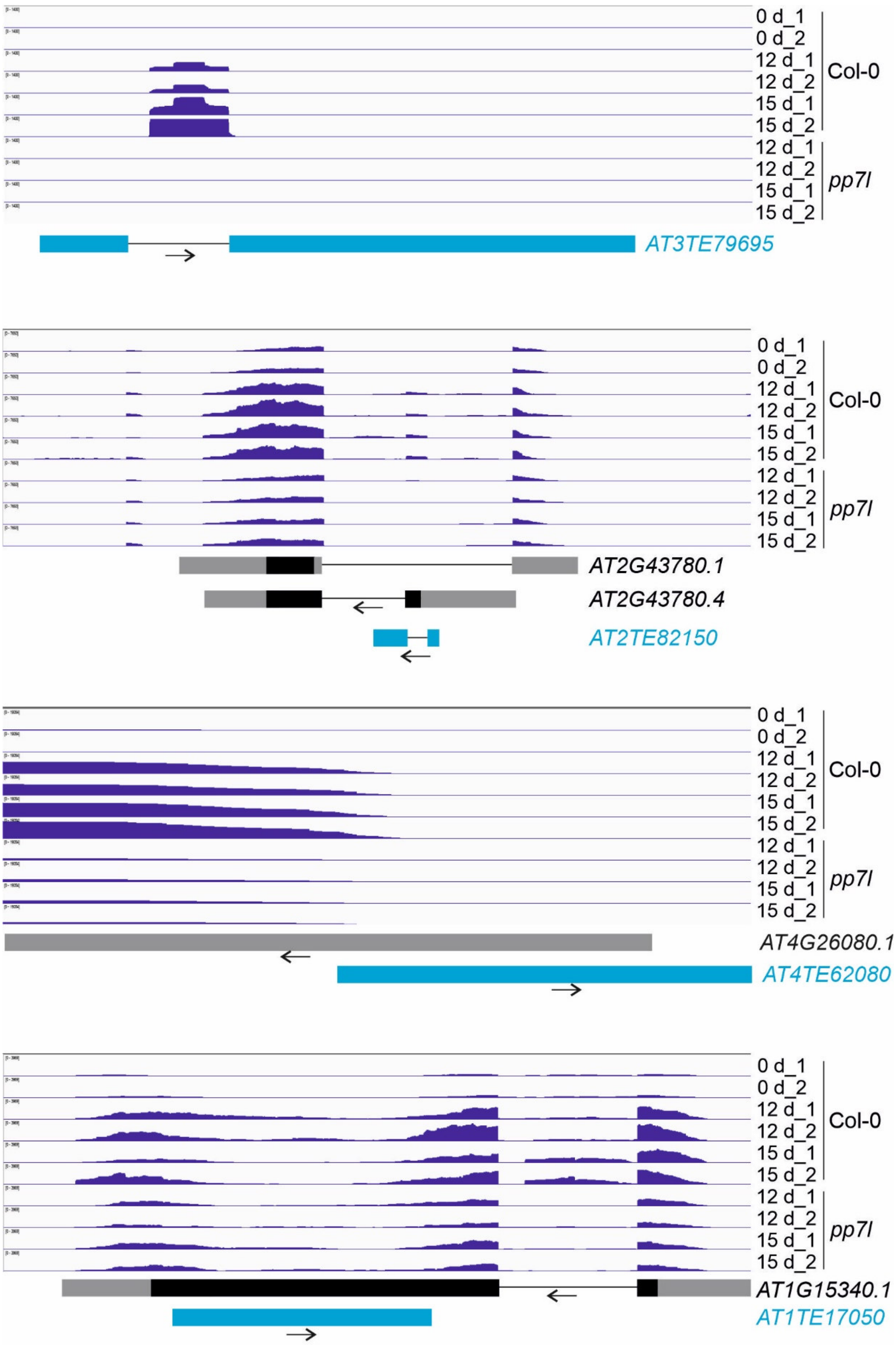


**Supplemental Figure S2.** Patterns of transcript accumulation covering regions of TEs. The normalized read depths of transcripts detected in Col-0 and *pp7l* plants under control and prolonged drought conditions were visualized with the Integrative Genomics Viewer (IGV). Transposable elements (blue boxes), exons (black boxes), introns (black lines) and the 5′- and 3′-UTRs (gray boxes) are shown.

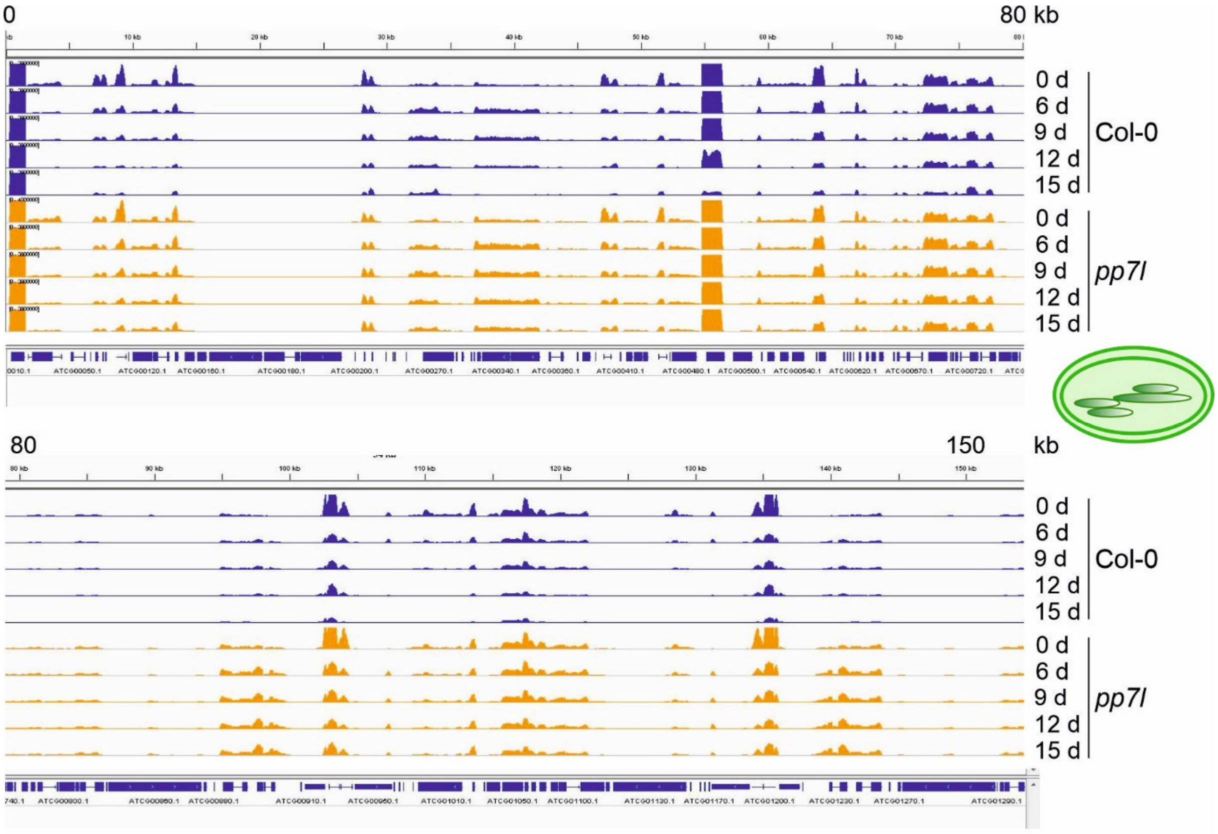


**Supplemental Figure S3.** Distribution of lncRNA-Seq reads across the chloroplast genome. The normalized read depths of transcripts detected in Col-0 and *pp7l* plants under control (0 d) and drought conditions were visualized with the Integrative Genomics Viewer (IGV).

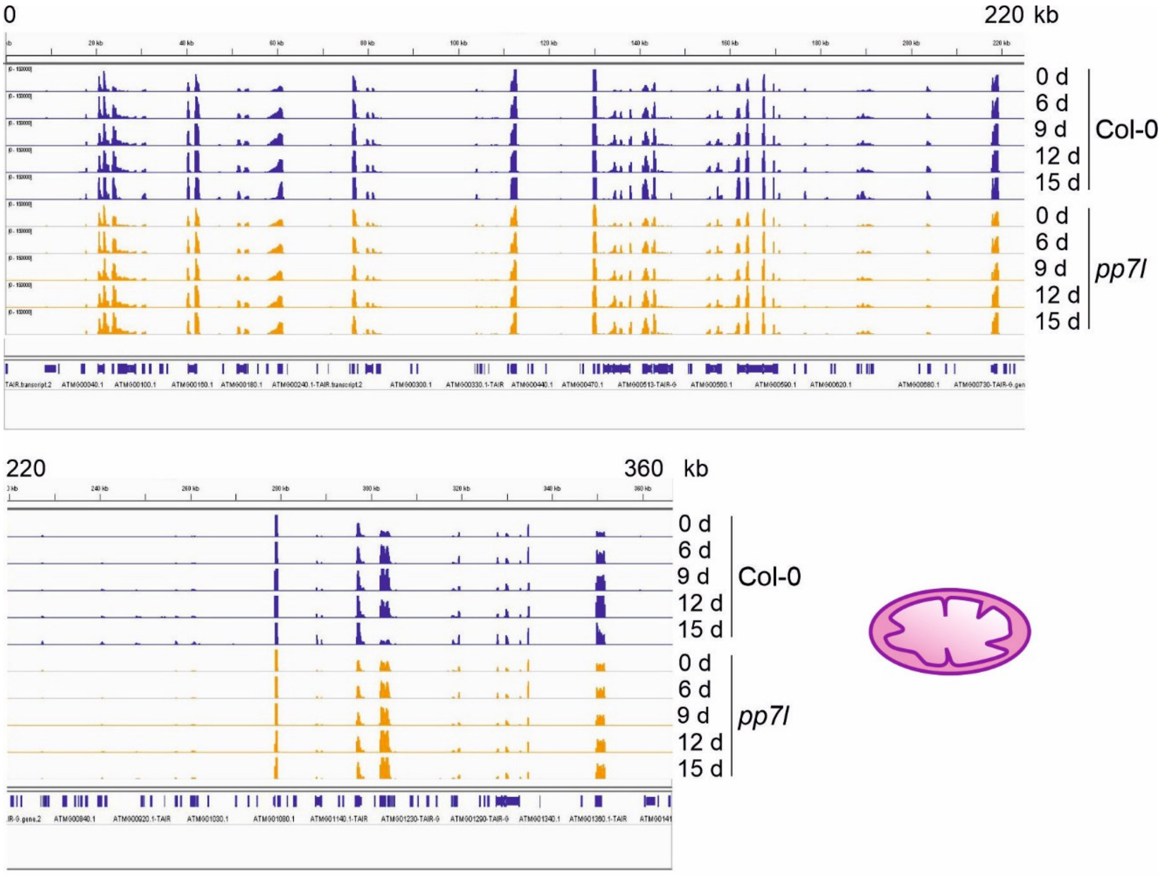


**Supplemental Figure S4.** Distribution of lncRNA-Seq reads across the mitochondrial genome. The normalized read depths of transcripts detected in Col-0 and *pp7l* plants under control (0 d) and drought conditions were visualized with the Integrative Genomics Viewer (IGV).

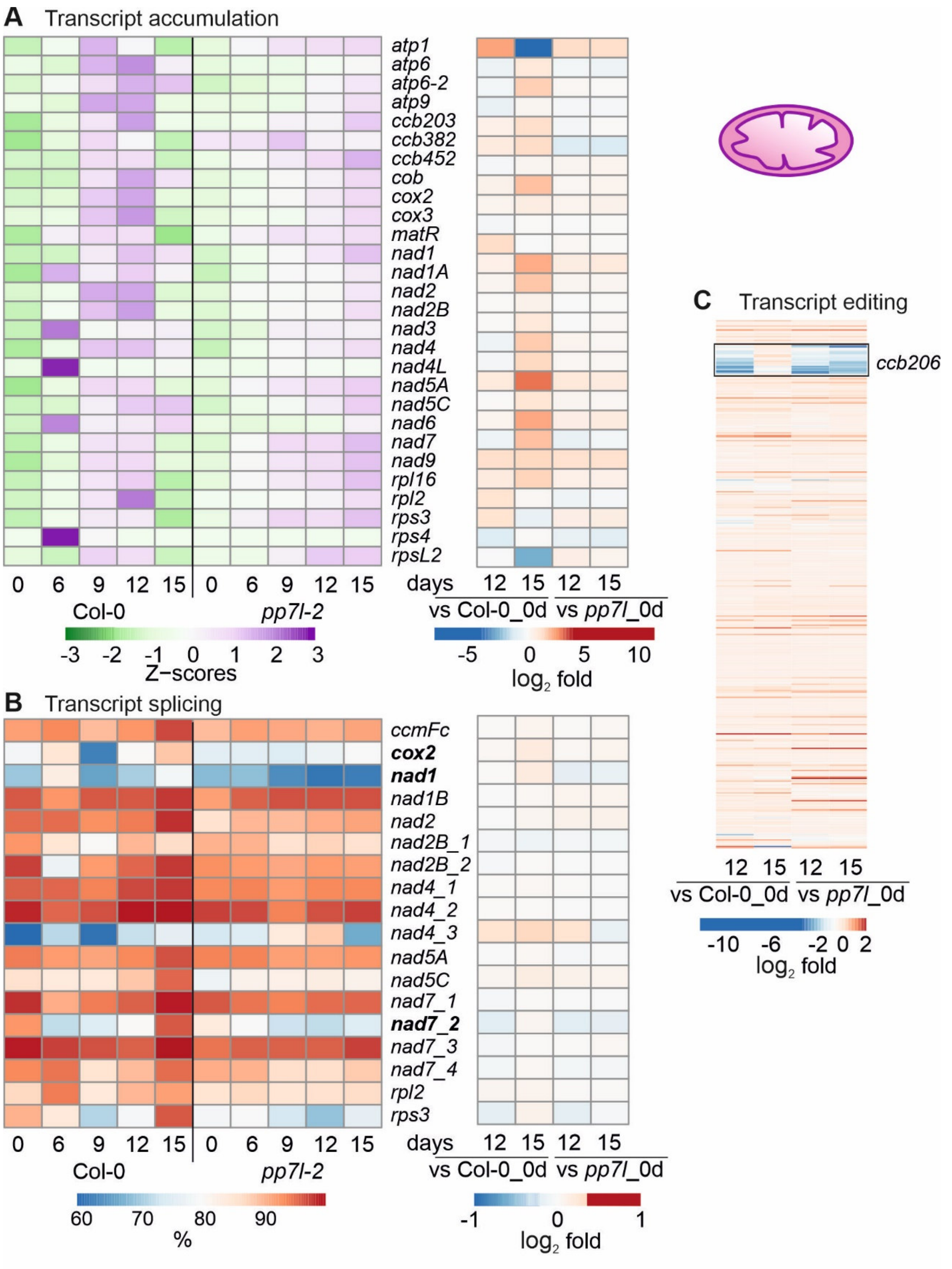


**Supplemental Figure S5.** Impact of drought stress on the accumulation, editing and splicing of mitochondrial transcripts. Plants and RNA were treated and analyzed, and data are depicted as described in the legend to Figure 3. **(A)** Heatmap illustrating chloroplast transcript accumulation (Z-scores) during the drought time-course. **(B)** Percentages of transcript splicing during the time course (left) and $\log_2$ fold changes after 12 and 15 days of drought stress compared to the control time point (right side). **(C)** $\text{Log}_2$ fold changes of transcript editing during the time course.

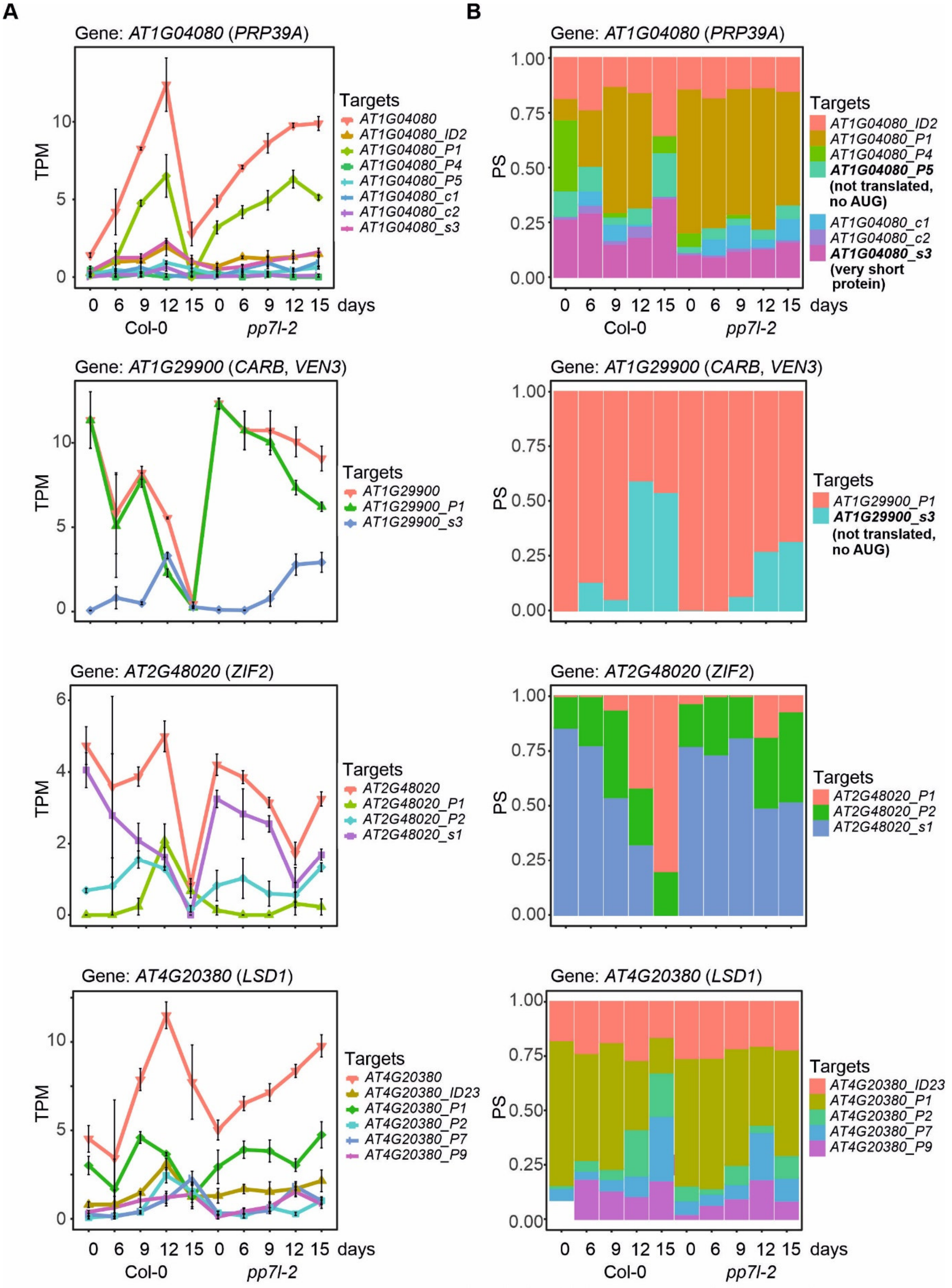


**Supplemental Figure S6.** Illustrations of further isoform switching (IS) events. Expression profiles of *PRP39A, CARB, ZIG2* and *LSD1* at the whole-gene level and at the level of detected transcript isoforms are shown. PS, percentage of expressed transcripts spliced; TPM, transcripts per million reads.

**Supplemental Table S9**. Primers used in this study.

| Atg number | Description | Primer sequence (from 5' to 3') |
|---|---|---|
| **Genotyping** | | |
| *AT5G10900* | *pp7l-1*-SALK_018295 LP | ATGCCGTCAACTTCAACAATC |
| | *pp7l-1*-SALK_018295 RP | CATTCTTGAAGCTAAGTGCGG |
| *AT5G10900* | *pp7l-2*-SALK_003071 RP | ATTGTTGAAGTTGACGGCATC |
| | *pp7l-2*-SALK_003071 LP | CCAGTTTCTTACCCTTGGGAC |
| | SALK LB | GTCCGCAATGTGTTATTAAGTTGTC |
| **qRT-PCR** | | |
| *AT5G10900* | *PP7L* | AGTCTTCAGTGCTTCAATGTTCTC |
| | | GATTTGATGTTGTAACCTTGGGAC |
| *AT1G77080* | *FLM-RT_F* | TCGCTGTTGTCGTCGTATCTGC |
| *AT1G77080* | *FLM-e1-e3-RT_R* | CGGCTTGAACAGCGCTTCTATCTC |
| *AT1G77080* | *FLM-e1-e2-RT_R* | CAATGATCTTGGAAATGTCGTCACCG |
| *AT1G77080* | *FLM-intr1-RT_R* | GAAGCTTCTATATGGAGAAAGTAA |

**In separate files:**

**Supplemental Table S1.** Sequencing depth of lncRNA-Seq experiments. Long non-coding RNA sequencing (lncRNA-Seq) was performed with RNA isolated from 3-week-old Col-0 (Col) and *pp7l* plants grown under optimal conditions (time-point 0 days; 0d) and after water had been withheld for 6, 9, 12 and 15 days, respectively (6d, 9d, 12d and 15d). Sequences were mapped and analyzed as described in Materials and Methods.

**Supplemental Table S2.** Genes differentially expressed under drought stress. Long non-coding RNA sequencing (lncRNA-Seq) was performed with RNA isolated from 3-week-old Col-0 (Col) and *pp7l* plants grown under optimal conditions (time-point 0 days; 0d) and after water had been withheld for 6, 9, 12 and 15 days, respectively (6d, 9d, 12d and 15d). Sequences were mapped and analyzed with the 3D RNA-Seq pipeline as described in Materials and Methods.

**Supplemental Table S3.** Drought-stress marker genes. Lists of genes whose transcripts are regulated in the same direction (up or down) after 9, 12 and 15 days of drought stress in our dataset as well as in Kim et al. (2017).

**Supplemental Table S4.** Gene Ontology (GO) analysis of the nine clusters listed in Figure 5. Biological process and cellular component categories according to DAVID (Huang da et al., 2009) and REVIGO (Supek et al., 2011).

**Supplemental Table S5.** Normalized read counts across regions containing transposable elements (TEs).

**Supplemental Table S6.** Abundance of chloroplast and mitochondrial transcripts. **A,** $Log_2$ fold changes of RNAs encoded by the chloroplast genome. Note that analysis of tRNAs was excluded. **B,** $Log_2$ fold changes of RNAs encoded by the mitochondrial genome.

**Supplemental Table S7.** List of differentially alternatively spliced (DAS) genes under drought stress.

**Supplemental Table S8.** List of genes that showed isoform switches (ISs) during drought stress.

### References


**Huang da, W., Sherman, B.T., and Lempicki, R.A.** (2009). Bioinformatics enrichment tools: paths toward the comprehensive functional analysis of large gene lists. Nucleic Acids Res **37,** 1-13.

**Supek, F., Bosnjak, M., Skunca, N., and Smuc, T.** (2011). REVIGO summarizes and visualizes long lists of gene ontology terms. PLoS One **6,** e21800.
